# Virus-free and live-cell visualizing SARS-CoV-2 cell entry for studies of neutralizing antibodies and compound inhibitors

**DOI:** 10.1101/2020.07.22.215236

**Authors:** Yali Zhang, Shaojuan Wang, Yangtao Wu, Wangheng Hou, Lunzhi Yuan, Chenguang Sheng, Juan Wang, Jianghui Ye, Qingbing Zheng, Jian Ma, Jingjing Xu, Min Wei, Zonglin Li, Sheng Nian, Hualong Xiong, Liang Zhang, Yang Shi, Baorong Fu, Jiali Cao, Chuanlai Yang, Zhiyong Li, Ting Yang, Lei Liu, Hai Yu, Jianda Hu, Shengxiang Ge, Yixin Chen, Tianying Zhang, Jun Zhang, Tong Cheng, Quan Yuan, Ningshao Xia

**Affiliations:** State Key Laboratory of Molecular Vaccinology and Molecular Diagnostics, School of Public Health & School of Life Sciences, Xiamen University, Xiamen 361102, Fujian, China; National Institute of Diagnostics and Vaccine Development in Infectious Diseases, School of Public Health & School of Life Sciences, Xiamen University, Xiamen 361102, Fujian, China; Shenzhen Key Laboratory of Pathogen and Immunity, National Clinical Research Center for Infectious Disease, Shenzhen Third People’s Hospital, Second Hospital Affiliated to Southern University of Science and Technology, Shenzhen 518112, Guangdong, China; Department of Hematology, Fujian Medical University Union Hospital, Fujian Provincial Key Laboratory on Hematology, Fujian Institute of Hematology, Fuzhou 350001, Fujian, China; The First Hospital of Xiamen University, Xiamen 361003, China

**Keywords:** SARS-CoV-2, coronavirus, spike glycoprotein, fluorescent protein, live-cell imaging, viral entry visualization, high-throughput screen, high-content analysis, entry inhibitor

## Abstract

The ongoing COVID-19 pandemic, caused by SARS-CoV-2 infection, has resulted in hundreds of thousands of deaths. Cellular entry of SARS-CoV-2, which is mediated by the viral spike protein and host ACE2 receptor, is an essential target for the development of vaccines, therapeutic antibodies, and drugs. Using a mammalian cell expression system, we generated a recombinant fluorescent protein (Gamillus)-fused SARS-CoV-2 spike trimer (STG) to probe the viral entry process. In ACE2-expressing cells, we found that the STG probe has excellent performance in the live-cell visualization of receptor binding, cellular uptake, and intracellular trafficking of SARS-CoV-2 under virus-free conditions. The new system allows quantitative analyses of the inhibition potentials and detailed influence of COVID-19-convalescent human plasmas, neutralizing antibodies and compounds, providing a versatile tool for high-throughput screening and phenotypic characterization of SARS-CoV-2 entry inhibitors. This approach may also be adapted to develop a viral entry visualization system for other viruses.

## Introduction

Since previous outbreaks of SARS-CoV-1 in 2002 and MERS-CoV in 2012, COVID-19 caused by SARS-CoV-2 infection has become pandemic ^1–4^. The development of therapeutic and preventative agents against SARS-CoV-2 infection is urgently needed. Viral cellular entry is the first step for the establishment of a productive viral infection^5^. Effective inhibition of viral entry is an important goal for the development of antiviral antibodies, vaccines, and drugs ^6–8^. The cell entry of SARS-CoV-2 is mediated by viral spike (S) glycoprotein and its interaction with the cellular ACE2 receptor ^9–14^. To date, a variety of approaches have been employed to develop prophylactic and therapeutic measures aimed at functional blockage of SARS-CoV-2 cell entry ^15–19^.

Current cell-based assays for study SARS-CoV-2 cell entry, using either authentic virus or spike-bearing pseudotyping virus^20–22^, require biosafety facilities and multistep experimental procedures and are time-consuming, which has greatly limited relevant studies, particularly high-throughput screening studies. With the goal of establishing an ideal system for high-throughput screening of SARS-CoV-2 entry inhibitors in virus-free conditions and facilitating the development of antibodies and vaccines, we developed a fluorescent SARS-CoV-2 entry probe that can be visualized and quantified via live-cell imaging. Using the novel probe, we established a one-step ultrafast assay for characterization of various SARS-CoV-2 entry inhibitors. The practical applicability of the new system was systematically evaluated by using human COVID19-convalescent plasmas, immunized mouse sera, monoclonal antibodies (mAbs) and compound inhibitors.

### Recombinant FP-fused spike proteins of coronaviruses

The constructs used to produce the recombinant FP-fused coronavirus spike probes contain the following elements: (i) an N-terminal signal peptide; (ii) a receptor-binding domain (RBD) or the S-ectodomain; (iii) a flexible-linker following green fluorescent protein (GFP); and (iiii) a T4-fibritin foldon (TFd) for trimerization the S-ectodomain (Figure 1A). The Gamillus (mGam) and mNeonGreen (mNG) were tested as the fused-GFP because mGam is acid-tolerant, which may enable fluorescent tracking when the probe is taken up into acidic cellular organelles ^23^, and mNG is the brightest GFP to our knowledge ^24^. We designated the RBD-based probes as RBG (mGam-fused) or RBN (mNG-fused) and designated the S-ectodomain trimer (ST)-based probes as STG (mGam-fused) and STN (mNG-fused). We expressed recombinant RBG proteins for the SARS-CoV-2, SARS-CoV-1, MERS, HKU1 and RaTG13 coronaviruses and STG and STN probes for SARS-CoV-2 in CHO cells (Figure 1B and Figure S1). Non-FP-fused SARS-CoV2-RBD and SARS-CoV2-ST proteins and a nontrimerized mGam-fused S-ectodomain (SARS-CoV2-SMG) were also produced. The molecular weights of the SARS-CoV2-STG and SARS-CoV2-STN were determined to be approximately 808-kd by size-exclusion chromatogram (Figure 1C, Figure S2A-B). Furthermore, Cryo-EM reconstructions of the SARS-CoV2-ST (Figure S2C) and SARS-CoV2-STN (Figure S2D) both demonstrated a typical trimeric structure ^9, 10^. The binding affinities of SARS-CoV2-STG and SARS-CoV2-RBG to human ACE2 (hACE2) were 18.2 nM and 30.4 nM (Figure 1D), respectively, which were similar to previously reported data for unfused proteins ^9, 10^. Together, C-terminal FP-fusion does not influence the structure and ACE2-binding capability of the RBD and S-ectodomain of SARS-CoV-2.

**Figure 1.**
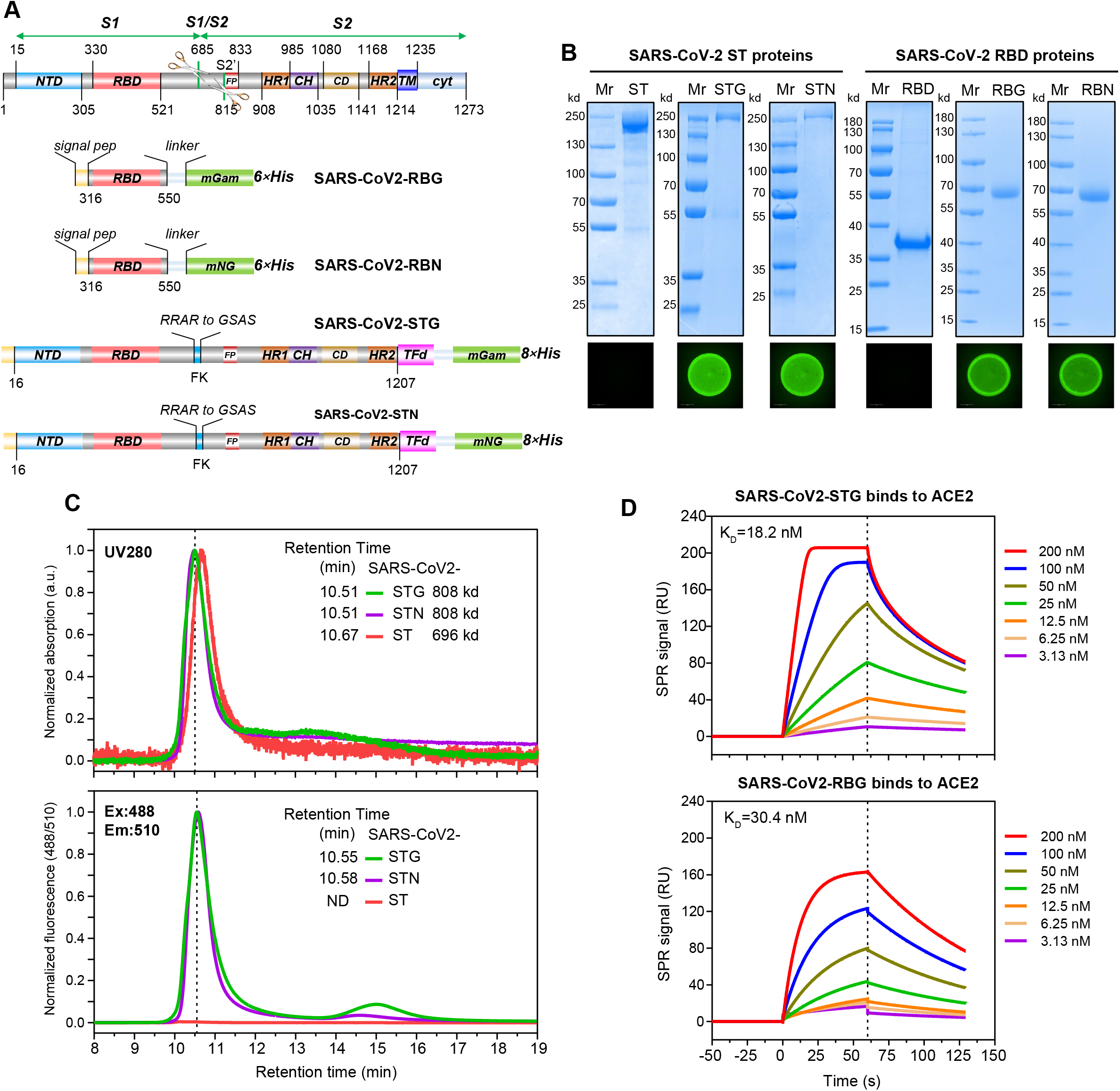
Generation and characterization of FP-fused SARS-CoV-2 S proteins. (A) Schematics of STG and RBG constructs. Functional domains are colored. NTD, N-terminal domain; RBD, receptor binding domain; FP, fusion peptide; HR1/2, heptad repeat 1/2; CH, central helix; TM, transmembrane domain; cyt, cytoplasmic tail; TFd, T4 fibritin trimerization motif; mGam, monomeric Gamillus; mNG, mNeonGreen. (B) SDS-PAGE and fluorescence analyses for purified ST-based and RBD-based SARS-CoV-2 S proteins. (C) Size-exclusion chromatogram (SEC) of the purified SARS-CoV2-ST, SARS-CoV2-STG and SARS-CoV2-STN. Data from UV280 detector (upper panel) and fluorescence detector (lower panel) from a G3000 HPLC Column were showed. The molecular weight of SARS-CoV2-STG (or SARS-CoV2-STN) was about 808 kd, which was calculated according to its elution time in referring to the standard curve of determining the molecular weight as shown in Figure S2A and S2B. (D) SPR sensorgrams showing the binding kinetics for SARS-CoV2-STG (upper panel) or SARS-CoV2-RBG (lower panel) with immobilized rACE2 (human). Colored lines represented a global fit of the data using a 1:1 binding model.

### Establishment of virus-free assays to visualize SARS-CoV-2 cell entry

We established hACE2-overexpressing cell lines using the ACE2hR and ACE2iRb3 constructs (Figure 2A). Cell-transfection with ACE2hR allowed hACE2-overexpression with nucleus visualization (H2B-mRuby3). ACE2iRb3 contains an ACE2-mRuby3 expressing cassette following an IRES-ligated H2B-iRFP670-2A-PuroR. Transfection with ACE2iRb3 simultaneously enabled fluorescent visualization of hACE2 (hACE2-mRuby3) and nucleus (H2B-iRFP670). Using these vectors, we developed three stable cell lines, namely, 293T-ACE2iRb3, 293T-ACE2hR and H1299-ACE2hR. As expected, hACE2 (or ACE2-mRuby3) was expressed at high levels, and the expression of TMPRSS2 (another critical factor for viral entry) ^12^ did not change in these cells (Figure 2B).

**Figure 2.**
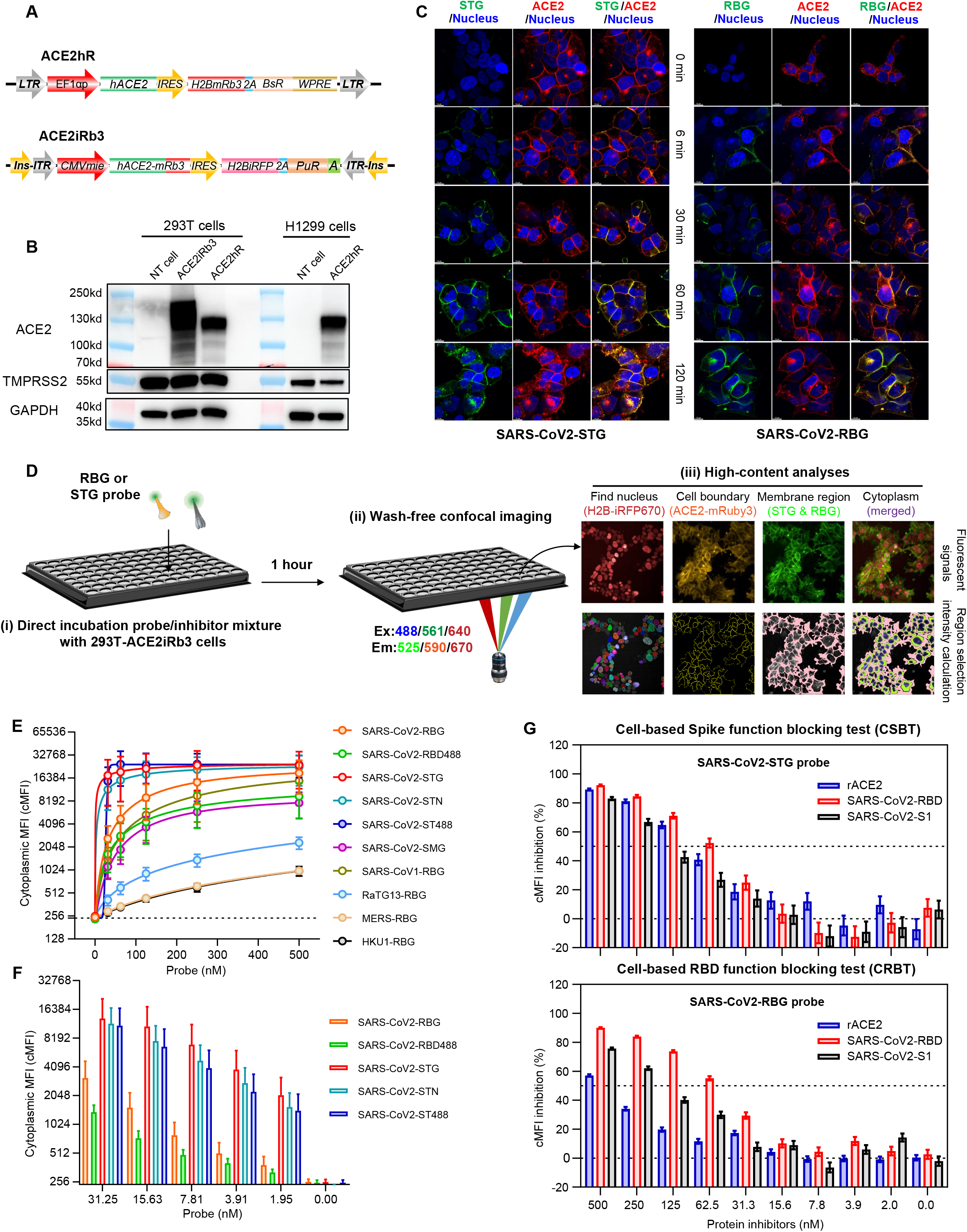
Establishment of the CSBT and CRBT assays. (A) Schematics of the constructs of ACE2hR and ACE2iRb3 for generations of ACE2-overexpressing cell lines. EF1αp, human EF-1 alpha promoter; hACE2, human ACE2; IRES, internal ribosome entry site; H2BmRb3, H2B-fused mRuby3; BsR, blasticidin S-resistance gene; 2A, P2A peptide; ins, insulator; hCMVmie, a modified CMV promoter derived from pEE12.4 vector; hACE2-mRb3, human ACE2 with C-terminal fusing of mRuby3; H2BiRFP, H2B-fused iRFP670; PuR, puromycin resistance gene. (B) Western blot analyses of expressions of ACE2 and TMPRSS2 in 293T and H1299 cells stably transfected with different constructs. NT cell, non-transfected cells. (C) Fluorescence confocal images of 293T-ACE2iRb3 cells incubated with SARS-CoV2-RBG and SARS-CoV2-STG for different times. The nucleus H2B-iRFP670 was pseudo-colored blue. The scale bar was 10 μm. (D) Schematic illustration of the procedures of cell-based high-content imaging assay using fluorescent RBG or STG viral entry sensors. (E) Dose-dependent fluorescence responses (cMFI) of various probes derived from different CoVs on 293T-ACE2iRb3 cells. SARS-CoV2-RBD488 was a dylight488-conjugated SARS-CoV2-RBD protein, and SARS-CoV2-ST488 was a dylight488-conjugated SARS-CoV2-ST protein. Each probe was tested at 500, 250, 125, 62.5, and 31.25 nM, respectively. (F) Comparisons of the fluorescence response (cMFI) of various SARS-CoV-2 probes on 293T-ACE2iRb3 cells. For panel E and F, cell images were obtained for 25 different views for each test, and the data were expressed as mean±SD. (G) Dose-dependent cMFI inhibition of recombinant ACE2, SARS-CoV2-RBD, and SARS-CoV2-S1 proteins for the binding and uptake of SARS-CoV2-STG (upper panel) and SARS-CoV2-RBG (lower panel). The experiments were performed following the procedure as described in panel D. The data were mean±SD. CSBT, cell-based spike function blocking test; CRBT, cell-based RBD function blocking test.

On 293T-ACE2iRb3 cells, both the SARS-CoV2-RBG and SARS-CoV2-STG probes showed effective-binding to the cells, as membrane-bound and hACE2-mRuby3-colocalized mGam signals were observed after a 6-min incubation with the cells (Figure 2C). Cytoplasmic mGam signals were detected in cells after incubation for 60-min or longer with the probes, particularly for SARS-CoV2-STG, suggesting that the recombinant probes can not only bind to the cell surface but also be taken up into the cells. The internalization of SARS-CoV2-STG was more evident than that of SARS-CoV2-RBG into 293T-ACE2iRb3 (Figure 2C). In live-cell dynamic tracking, more internalized mGam signals was observed for SARS-CoV2-STG than SARS-CoV2-RBG (Figure S3A). For mGam signals, the internalized fluorescence ratio (IFR, Figure S3B) and the internalized vehicle numbers (IVNs, Figure S3D) of STG-treated cells were both significantly higher than those of RBG-treated cells approximately 30-min after probe-cell incubation. In contrast, no significant difference was noted in hACE2-mRuby3 internalization in the presence or absence of probes (Figure S3C).

Using 293T-ACE2iRb3 cells, we established a cell-based assay mimicking SARS-CoV-2 cell entry based on recombinant probes. It was a one-step wash-free assay (Figure 2D). After 1-hour cell-probe incubation, the cells were directly imaged by using a fully automatic high-content screening (HCS) system in confocal mode. For quantitative measurements, the H2B-iRFP670 were used to identify the nucleus, and the ACE2-mRuby3 were used to determine the cell boundary. Based on the detected nucleus and cell outlines, the green fluorescence intensities on the cell membrane and in the cytoplasmic region of each cell could be measured (Figure 2D). Generally, we used the mean fluorescence intensity (MFI) in the cytoplasmic region (cMFI) as an index of the amounts of the cell-bound and internalized probes. As the spikes of MRES-CoV and HKU1-CoV do not interact with hACE2, the signals of MERS-RBG and HKU1-RBG on 293T-ACE2iRb3 were nonspecific background (Figure 2E). RaTG13-RBG showed a detectable and dose-dependent cMFI, but the value was significantly lower than those of the probes of SARS-CoV-1 and SARS-CoV-2. Compared to SARS-CoV1-RBG, SARS-CoV2-RBG showed slightly stronger signals, possibly due to its higher binding affinity. For SARS-CoV-2, the cMFI of SARS-CoV2-STG, SARS-CoV2-STN and SARS-CoV2-ST488 were significantly higher than that of SARS-CoV2-RBG and SARS-CoV2-RBD488, and also stronger than SARS-CoV2-SMG. The dylight488-labeled SARS-CoV2-RBD488 probe presented a weaker signal than SARS-CoV2-RBG, suggesting that the NH2-dye modification at some amino acids of the RBD may interfere with its interaction with hACE2. Moreover, mGam-fused probes showed better performance than dylight488-labeled or mNG-fused probes (Figure 2F). At a concentration below than 10 nM, the cMFI of SARS-CoV2-STG was approximately 10-fold higher than that of SARS-CoV2-RBG.

Based on the visualization system, we developed cell-based HCS assays for analyzing the blocking potencies of SARS-CoV-2 entry inhibitors, designated CSBT (using SARS-CoV2-STG) and CRBT (using SARS-CoV2-STG), respectively. The proteins of hACE2-Fc (rACE2), SARS-CoV2-RBD and SARS-CoV2-S1 were employed for inhibition assessments following the procedure described in Figure 2D. As expected, all three proteins exhibited dose-dependent cMFI inhibition in both assays (Figure 2G). The Z’-factor coefficients of the CSBT and CRBT were both determined to be over 0.7 (Figure S3E), which demonstrated their robustness and reproducibility.

### Detecting entry-blocking antibodies in COVID-19-convalescent human plasmas by CSBT and CRBT

Recent studies have suggested that convalescent plasma may be beneficial in COVID-19 treatments ^25, 26^. Neutralization antibodies (NAbs) in convalescent plasmas may be essential in suppressing viruses ^27^. However, a rapid method for determining the neutralization antibody titer (NAT) of human plasma is still absent. We evaluated the feasibility of the CSBT and CRBT determined entry-blocking antibody titers as NAT surrogates in 32 COVID-19-convalescent human plasmas (Table S1). Compared with samples from healthy donors (n=40), all COVID-19-convalescent plasmas showed significant cMFI inhibition on CSBT assay, whereas only 12 samples (37.5%) had detectable CRBT activity (Figure 3A). For quantitative analysis, two-fold serial dilution tests were further performed to determine the CSBT and CRBT titers (Figure 3B). Moreover, the titers of total antibodies (TAb), IgG, IgM, and lentiviral-pseudotyping-particles (LVpp) based NAT (LVppNAT) against SARS-CoV-2 were also measured for comparisons (Figure 3C). Among the antibody titers derived from various assays, the CSBT titer showed the best correlation with LVppNAT (Figure 3D and Table S2, r=0.832, p<0.001), and it also well correlated (r=0.959, p<0.001, Figure 3D) with the neutralization activity against authentic SARS-CoV-2 virus in 12 representative samples (Table S3). Together, the CSBT-determined entry-blocking antibody titer is a good NAb surrogate of convalescent plasmas.

**Figure 3.**
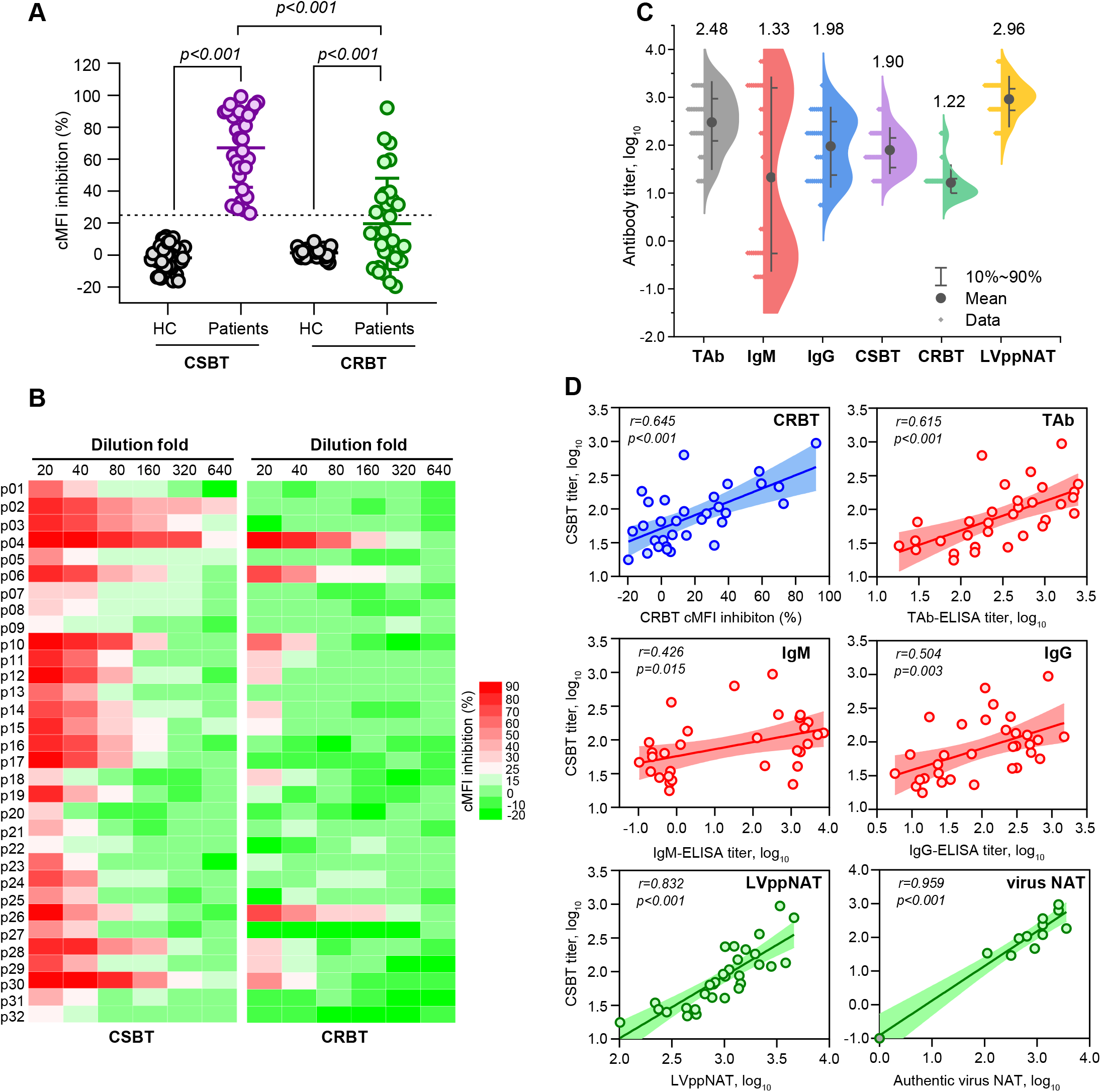
Evaluation of neutralization potential of human plasmas from convalescent COVID-19 patients by CSBT and CRBT assays. (A) Comparisons of cMFI inhibitions on CSBT and CRBT assays between plasma samples from convalescent COVID-19 patients and healthy control (HC) subjects. The cMFI inhibition (%) at 1:20 dilution was plotted at the left Y-axis. The cutoff values for CSBT and CRBT were inhibition of 25% (median HC value +3.3×SD) on cMFI at 1:20 dilution. (B) Heatmaps showing CSBT and CRBT effects of two-fold serial dilutions of 32 plasmas from convalescent COVID-19 patients. (C) Distributions of the levels of TAb, IgM, IgG, CSBT, CRBT and LVppNAT of convalescent plasma samples. The numbers indicated the average titers at log10. The titers of Ab, IgM, and IgG were expressed as relative S/CO values determined by serial dilution measurements of each sample (maximum reactive dilution fold multiplied by S/CO). The CRBT and CSBT titers were expresses at ID25, whereas the LVppNAT was expressed as ID50. (D) Correlation analyses between the CSBT titer and the CRBT efficiency (at 1:20 dilution), the TAb titer, the IgM titer, the IgG titer, the LVppNAT and the NAT against authentic SARS-CoV-2 virus among convalescent plasmas. The correlation of CSBT titer and neutralization activity against authentic SARS-CoV-2 virus in 12 representative samples (included 11 convalescent COVID-19 plasmas and 1 control sample).

### Functional phenotyping of mouse anti-spike antibodies by CSBT and CRBT

Serum samples from mice immunized with the SARS-CoV2-RBD, SARS-CoV2-S1 and SARS-CoV2-S2 were collected for LVppNAT, CSBT and CRBT measurements. The SARS-CoV2-RBD and SARS-CoV2-S1 immunizations resulted in potent and comparable serum LVppNAT (Figure S4A), whereas SARS-CoV2-S2 raised little NAbs. The CSBT (Figure S4B) and CRBT (Figure S4C) assays also exhibited similar results to LVppNAT measurements. The ID50 correlation coefficient was 0.989 (p<0.001) between CSBT and LVppNAT and 0.925 (p<0.001) between CRBT and LVppNAT.

Using RBD-immunized mice, we developed 18 mAbs via RBD-ELISA screening following cell-based functional evaluations (as illustrated Figure 4A). These mAbs did not display much difference in ELISA-binding to SARS-CoV2-RBD, but 2 of them (8H6 and 15A9) showed significantly decreased ELISA-binding activities to SARS-CoV2-ST (Figure S5A). Based on epitope-binning assays using a cross-competitive ELISA, the mAbs could be divided into six groups (Figure S5B). All mAbs showed detectable but varied surface plasmon resonance (SPR) affinity (0.004-131 nM, Figure S6) to SARS-CoV2-RBD. Quantitative measurements of CSBT, CRBT and LVppNAT for the mAbs were further performed (Figure 4B to C, Figure S7A to C). Half of the mAbs exhibited high-to-moderate CSBT blocking potencies (IC50<30 nM), whereas the remaining ones showed low-to-no CSBT activities (Figure 4B). In comparisons of the dose-dependent cMFI inhibitions against SARS-CoV2-STG, SARS-CoV2-ST488, SARS-CoV2-RBG, and SARS-CoV1-RBG (Figure 4C), the profiles of most mAbs against SARS-CoV2-STG and SARS-CoV2-ST488 were similar, but the activities of 2B4 and 34B4 were dramatically decreased with SARS-CoV2-ST488 compared to SARS-CoV2-STG, suggesting that dye-conjugation may modify the epitopes of the two mAbs and hinder their bindings. Notably, seven mAbs exhibited striking enhancement at some dosage in the CRBT assays (Figure 4C and Figure S7B), but neither enhancement was noted in the CSBT nor LVppNAT tests. SPR analyses demonstrated that the Fabs of two representative CRBT-enhancing mAbs, 53G2 and 8H6, also showed a dose-dependent promoting effect on the RBD-ACE2 binding, whereas the Fabs of two CRBT-blocking mAbs (36H6 and 2B4) exhibited a dose-dependent reduction in the RBD-ACE2 interaction (Figure S8). Together, the CRBT-enhancing effects of these mAbs may be caused by the antibody-induced RBD conformation changes associated with increases in ACE2-binding capacity.

**Figure 4.**
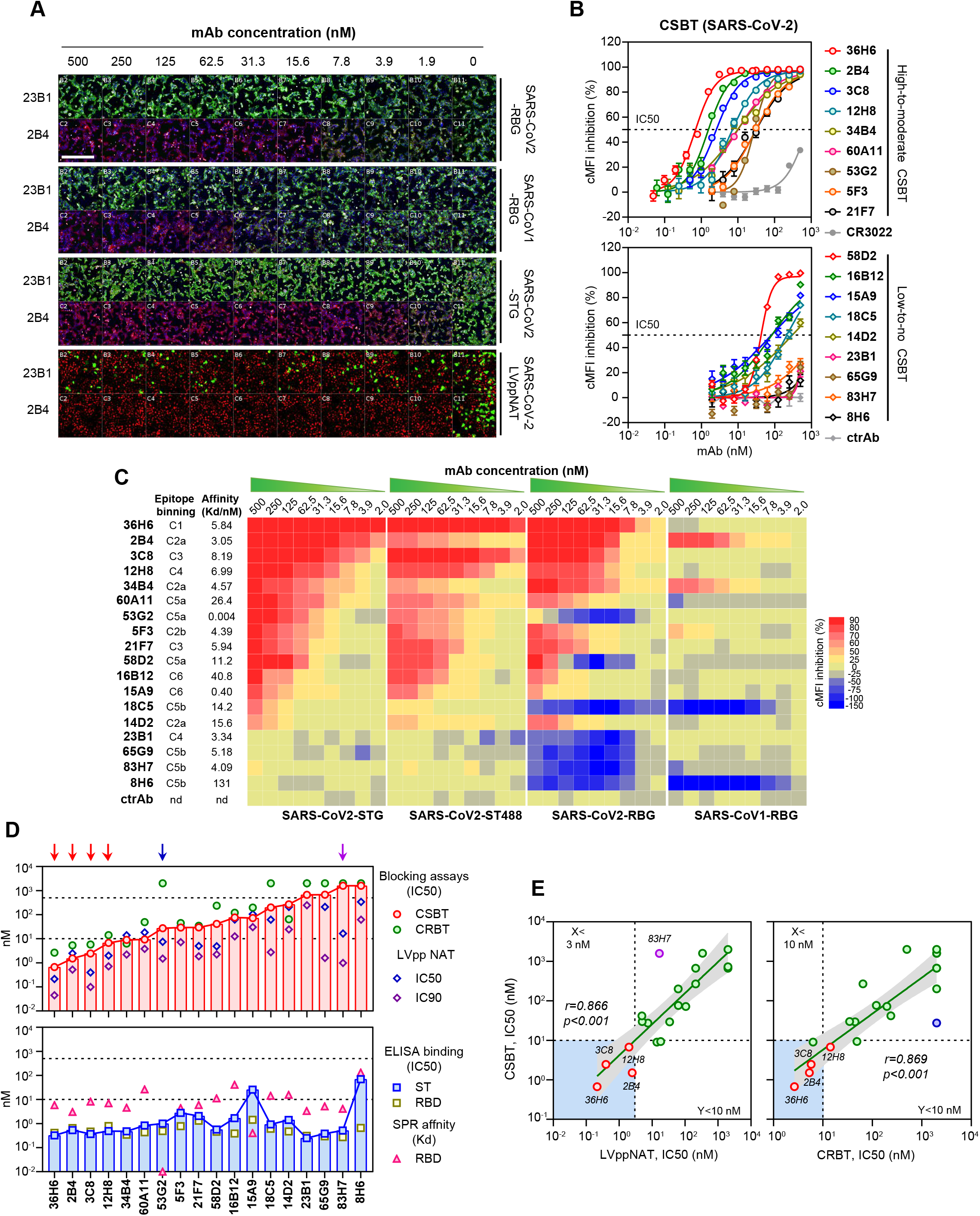
Phenotypic characterization of mAbs against by the CSBT and CRBT assays. (A) Fluorescence images for evaluations of inhibition effects of two representative mAbs (23B1 and 2B4) in tests of CRBT (SARS-CoV2-RBG and SARS-CoV1-RBG), CSBT (SARS-CoV2-STG) and LVppNAT (SARS-CoV-2). Scale bar, 500 μm. (B) CSBT titrations of mAbs to determine their inhibitory activities in blocking the SARS-CoV2-STG internalization. (C) Heatmaps showing dose-dependent inhibitory effects of mAbs on cell-based functional blocking tests using the probes of SARS-CoV2-STG, SARS-CoV2-ST488, SARS-CoV2-RBG and SARS-CoV1-RBG. The affinity data of mAbs to SARS-CoV2-RBD and the epitope binning cluster of the mAbs were shown on the left side of the pictures. (D) Comparison of potencies of mAbs determined by various cell-based functional assays (CSBT, CRBT, and LVppNAT) and ELISA or SPR-based binding assays. Red arrows indicated 4 mAbs (36H6, 2B4, 3C8 and 12H8) with the strongest CSBT blocking activities (IC50<10 nM) and potent neutralization activities (IC90<3 nM). A blue arrow indicated the 53G2 mAb which had CSBT but no CRBT activity. A purple arrow indicated the 83H7 mAb which had neutralization activity but showed neither CSBT nor CRBT inhibition. (E) Correlation between the CSBT-IC50 and the LVppNAT-IC90 (left panel) or CRBT-IC50 (right panel) of mAbs involved in this study. The 36H6, 2B4, 3C8 and 12H8 mAbs showing an LVppNAT-IC90 <3 nM and a CSB-IC50 <10 nM were plotted as distinct red dots. The 83H7 mAb was plotted as a purple dot in left panel, and the 53G2 mAb was plotted as a blue dot in right panel.

The functional potencies of mAbs determined by various assays are summarized in Figure 4D and Table S4. The CSBT-IC50 values of the mAbs showed the good correlation with their LVppNAT IC90 (r=0.866, p<0.001, Figure 4E) or IC50 (r=0.750, p<0.001, Figure S9) values, and were also well correlated with their CRBT-IC50 values (r=0.869, p<0.001, Figure 4E). However, a 53G2 mAb presented CSBT activity but no inhibition in the CRBT assay, suggesting that its CSBT activity is independent of the direct blocking of the RBD-ACE2 interaction (Figure 4D). An 83H7 mAb with moderate LVppNAT activity but showed neither CSBT nor CRBT inhibition (Figure 4D), suggesting it may act through different mechanisms to achieve neutralization. No significant relationship was noted between the ELISA-or SPR-determined protein-binding activities and neutralization potencies of the mAbs (Figure S9).

According to the SPR (Figure S10) and CRBT analyses using SARS-CoV1-RBG (Figure 4C), the 2B4, 34B4, 5F3, 18C5, and 8H6 mAbs showed cross-reactivity to SARS-CoV-1 and RaTG13-CoV. However, only the 2B4 mAb had neutralization activity in SARS-CoV-1 LVppNAT measurements (Figure S7D). Epitope-binning assay (Figure S5B) suggested that 2B4, 34B4, 5F3 and 14D2 possibly share an overlapping-epitope (cluster C2), and 18C5, 8H6, 83H7 and 65G9 may bind to another similar epitope (mAb cluster C5b). As 2B4 showed comparable LVppNAT potencies against both SARS-CoV-1 and SARS-CoV-2, it may recognize a cross-neutralization epitope. The 36H6 mAb, which recognizes a unique epitope that differs from other mAbs (mAb cluster C1, Figure S5B), presented the best performance in LVppNAT, CSBT, and CRBT assays but did not show any cross-reactivity with SARS-CoV-1 or RaTG13. Both 36H6 and 2B4 have neutralization activities against the authentic SARS-CoV-2 virus (Figure S7E), and the 36H6 exhibited superior neutralization activity with an IC50 of 0.079 nM (11.9 ng/mL).

### Characterization of the neutralization mechanisms of mAbs by STG-based viral entry visualizing system

The 83H7 mAb showed SARS-CoV-2 neutralization activity against both pseudoparticles (IC50=0.99 nM) and the authentic virus (IC50=13.02 nM) but had neither CSBT nor CRBT activity. We speculated that this mAb may inhibit the SARS-CoV-2 via an intracellular neutralization pathway ^28^. To validate this, we prepared dylight633-labeled mAbs (Ab633) of 36H6, 53G2, 83H7, and 8H6 and an irrelevant mAb (ctrAb) for dual-visualizing tracking. Among them, 36H6, 53G2 and 8H6 served as controls that had strong, moderate and weak/no activity for both CSBT and neutralization, respectively. In 293T-ACE2iRb3 cells simultaneously incubated with STG and Ab633, we performed time-serial live-cell imaging analyses. To characterize the influence of these mAbs on SARS-CoV2-STG internalization, the dynamic changes of the STG-IVNs, the STG-IVpMFI (the peak MFI of internalized vesicles), the Ab633-IVNs, the Ab633-IVpMFI, and the percentage of STG/Ab633-colocalized internalized vesicles, and the STG-IVA were calculated (Figure 5). As expected, 36H6 completely obstructed STG internalization (p<0.001), 53G2 showed significant but incomplete inhibition (p=0.002), and 8H6 presented little/no influence on STG internalization (p=0.70). Compared to ctrAb, the 83H7 showed no significant STG-IVNs reduction (p=0.31, Figure 5A) but increased the STG-IVpMFI (p<0.001, Figure 5B). On the other hand, the 83H7 group exhibited higher Ab633-IVNs (p<0.001, Figure 5C) and Ab633-IVpMFI (p<0.001, Figure 5D). The STG/Ab633 colocalization (p<0.001, Figure 5E) and the STG-IVA (p<0.001, Figure 5F) in the 83H7 group were also significantly higher and larger, respectively, than those in the other groups. Representative images at 5-hour post STG/Ab633-cell incubation, as shown in Figure 5G, further confirmed these findings. These results demonstrated that 83H7 could efficiently enter cells in the presence of the SARS-CoV-2 S protein. The enlarged internalized STG vesicles in associating with 83H7 suggested that the mAb may induce aggregation and disturb intercellular function of S protein, which may contribute to its neutralization activity.

**Figure 5.**
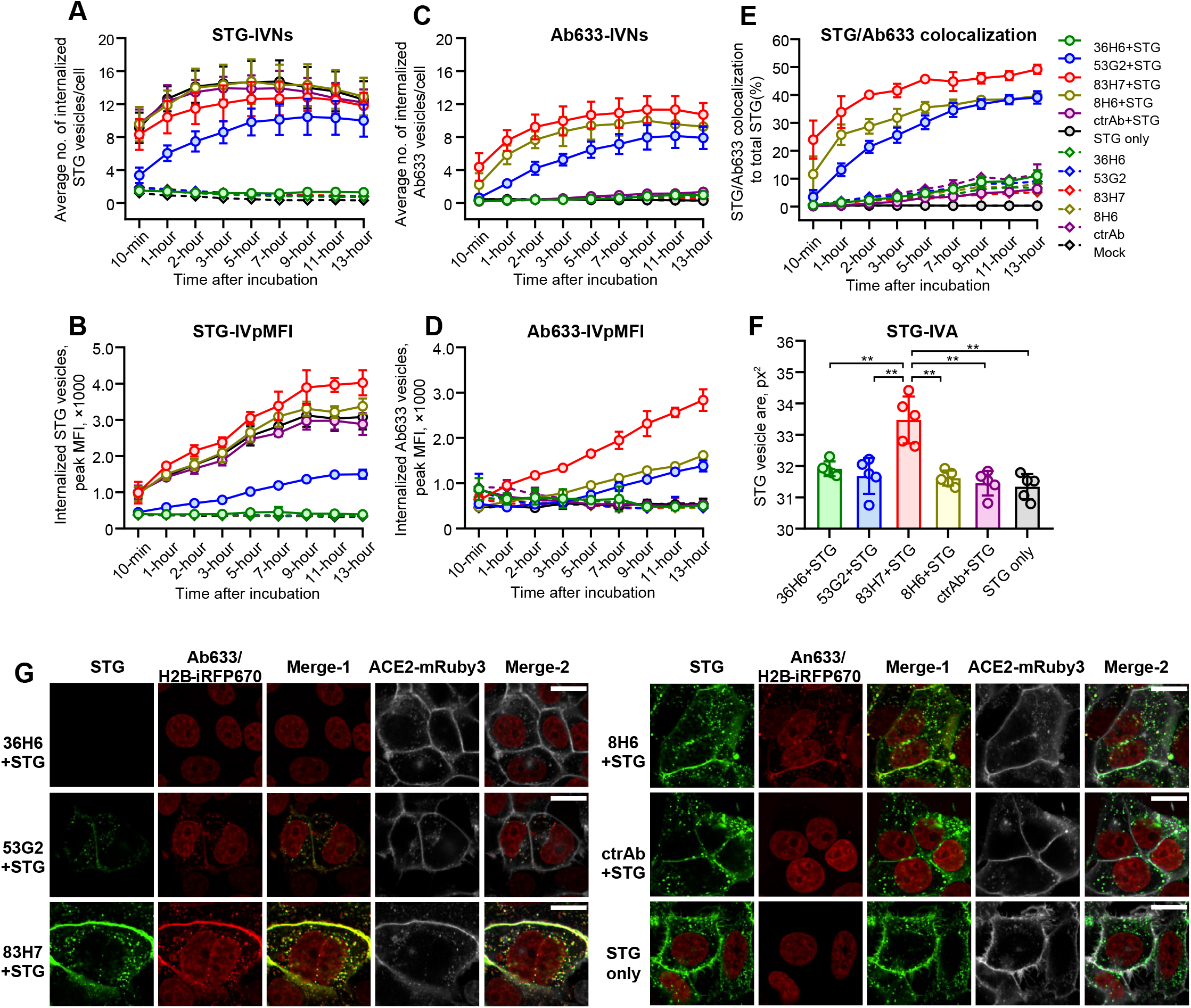
The 83H7 mAb inhibits SARS-CoV-2 via the intracellular neutralization pathway. The 293T-ACE2iRb3 cells were incubated with 20 nM of dylight633-labeled mAbs (Ab633) of 36H6, 53G2, 83H7, and 8H6 and an irrelevant control antibody (ctrAb), in the presence or absence of STG (2.5 nM). Live-cell fluorescence image dynamically tracked using a 63x water immersion objective. Five replicate wells were measured for each group, and 16 fields of each well were imaged. Time-series (at 10-min, 1-hour, 2-hour, 3-hour, 5-hour, 7-hour, 9-hour, 11-hour, and 13-hour) analyses of the STG-IVNs (A), STG-IVpMFI (B), Ab633-IVNs (C), Ab633-IVpMFI (D) and the percentage of STG/Ab633 colocalized vesicles to total internalized STG vesicles (E). IVNs, average internalized vesicle numbers; IVpMFI, the average peak MFI of internalized vesicles. (F) Comparisons of the STG-IVA of the internalized STG vesicles among groups co-incubated with various mAbs at 5-hour post-incubation. ** indicates p<0.01; IVA, average area (px^2^) of internalized STG vesicles. (G) Confocal images of STG (green channel), Ab633 (red channel), and ACE2-mRuby3 (white channel) in 293T-ACE2iRb3 cells at 5-hour post STG/Ab633 co-incubation. Scale bar, 20 μm.

### Visualization of the influence of compounds on SARS-CoV-2 cell entry

Previous studies suggested that the SARS-CoV-2 gains entry into cells via endocytosis. In this study, 11 inhibitors targeting various processes of endocytosis and endosome maturation were evaluated (Figure 6A). In SARS-CoV-2 LVpp infection tests (Figure 6B), the micropinocytosis inhibitor cytochalasin-D (CytD) and the clathrin-dependent endocytosis (CME) inhibitors dynasore and dansylcadaverine (MDC) showed dose-dependent inhibition with high-micromolar IC50 values. In contrast, neither of the caveolae-mediated endocytosis inhibitors nystatin and filipin inhibited viral infection. These results suggested that micropinocytosis and CME, instead of caveolae-mediated endocytosis, are involved in SARS-CoV-2 cell entry. Moreover, apilimod, a phosphoinositide 5-kinase (PIKfyve) inhibitor^29^, showed a low-nanomolar IC50. Similar effects were observed for two other PIKfyve inhibitors (YM201636 and APY0201). The acidification inhibitor bafilomycin A1 (Baf.A1) or the TPC2 inhibitor tetrandrine^30^ also significantly diminished viral infection.

**Figure 6.**
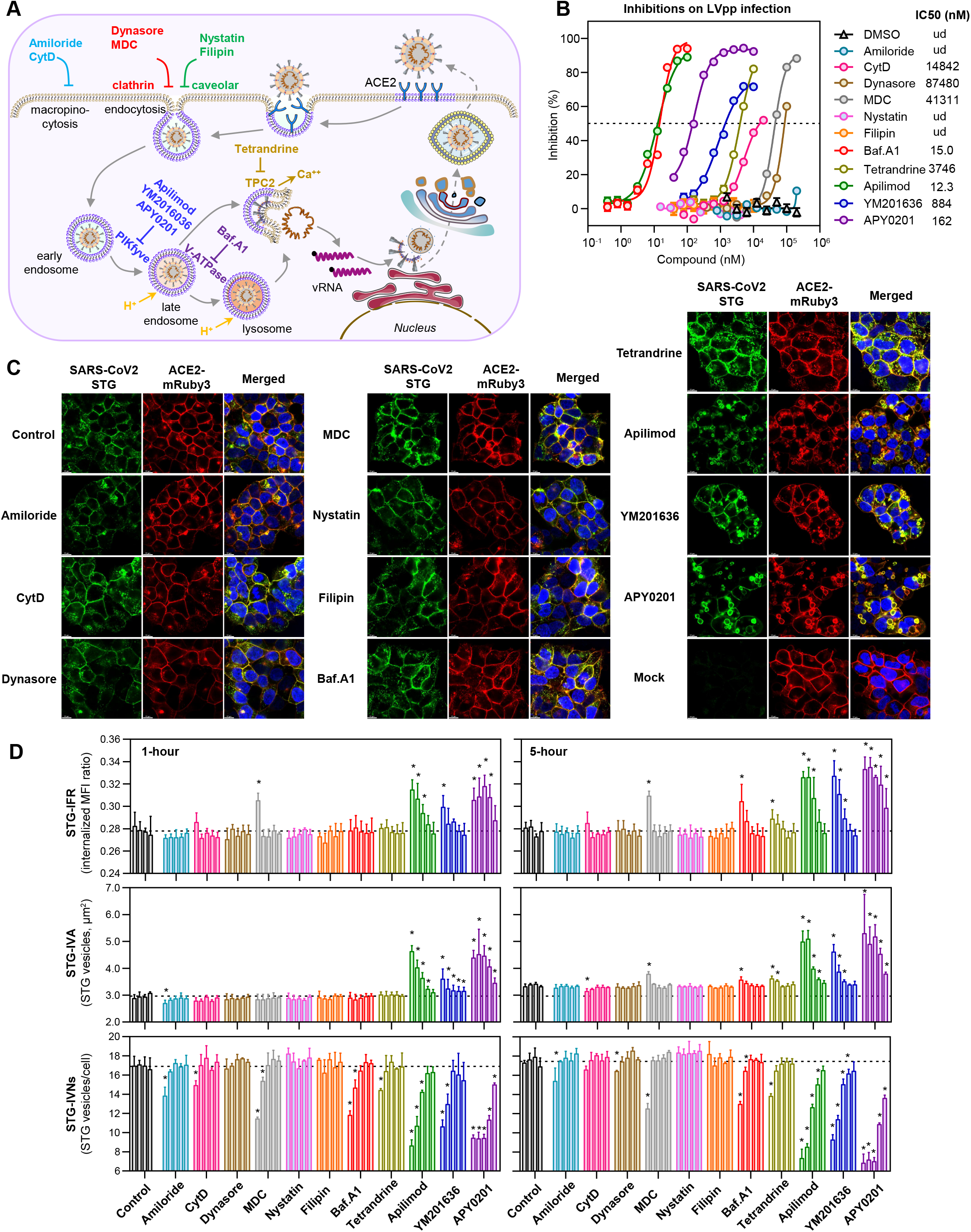
Detection of compound-induced influence on SARS-CoV-2 S-mediated cellular entry. (A) Schematic summary of the possible mechanisms of 11 compound inhibitors involved in the study. CytD, cytochalasin D; MDC, dansylcadaverine; Baf.A1, bafilomycin A1; vRNA, viral RNA. (B) Dose-dependent inhibitions of 11 compounds against SARS-CoV-2 LVpp infection on H1299-ACE2hR cells. All compounds were tested in a 2-fold dilution series, and the initial drug concentrations were begun at their maximal non-cytotoxic concentrations. The initial concentrations were 200 μM for amiloride, MDC and DMSO (as a solvent control); 100 μM for dynasore; 10 μM for filipin, APY0201, YM201636 and tetrandrine; 4 μM for nystatin; 100 nM for Baf.A1 and apilimod. ND, not detected. (C) Confocal images of STG (green channel), ACE2-mRuby3 (red channel), and nucleus (blue channel) in 293T-ACE2iRb3 cells at 5-hour post STG incubation. The cells were pretreated with compounds for 1-hour before STG loading. These pictures were obtained by using Leica gSTED confocal microscopy on cells treated with compounds at their respective initial concentrations as above-mentioned. Scale bar, 10 μm. (D) Quantitative analysis of the influence of entry inhibitors on STG internalization. Dose-dependent influence of various compounds on STG internalization characteristics on 293T-ACE2iRb3 cells at 1-hour (left panels) and 5-hour (right panels) after incubation. All compounds were tested in a 4-fold dilution series (4 gradients for DMSO control, and 5 gradients for others), and the initial drug concentrations were identical with as (B). Three replicate wells were measured for each group, and 16 fields of each well were imaged. For each compound, 5 colored bars from left-to-right orderly displayed the values measured from cells treated with 4-fold serial high-to-low concentrations of compounds. STG-IFR, internalized STG fluorescence intensity ratio; STG-IVA, average area (μm^2^) of internalized STG vesicles; STG-IVNs, average numbers of internalized STG vesicles per cell; *, p<0.05.

In STG-visualization system, the cMFI measurements following the CSBT procedure showed only a slight reduction in cells treated with a high dose of Amiloride, dynasore, apilimod, and APY0201 (Figure S11A). Notably, confocal-images revealed that the STG, colocalizing with internalized ACE2-mRuby3, were trapped on enlarged cytoplasmic vacuoles induced by PIKfyve inhibitors (Figure 6C), and most of these vacuoles were not stained by pH-dependent LysoView633 dyes, suggesting an abnormal pH-status (Figure S11B). In addition, tetrandrine or Baf.A1 also caused marked reductions in colocalization of internalized STG and LysoView633 stain-signals (Figure S11B), suggested that the two compounds also disturbed STG intracellular trafficking. For quantitative characterizations of the compound-induced influence on STG-internalization, the IVNs, IVA and IFR for cells treated with different concentrations of compounds at 1-hour and 5-hour post probe-loading were calculated. Apparently, compounds with infection-inhibitory effects correspondingly induced IVNs reduction or the increase of IVA or IFR (Figure 6D). The SARS-CoV-2 LVpp infection efficiencies were positively correlated with the IVNs and were negatively correlated with the IVA and IFR parameters (Figure S12). Overall, the 1-hour IVNs showed the best correlation with LVpp infection efficiencies (r=0.870, p<0.001). These results demonstrated the practical applicability of the STG-visualization system to screen and characterize compound inhibitors against SARS-CoV-2 cell entry.

## Discussion

Together, we established a versatile visualization system enables live-cell visualization of cellular binding, uptake, and intracellular trafficking of SARS-CoV-2 in virus-free conditions. The innovative system have several advantages over traditional technology: (i) using a recombinant FP-fused SARS-CoV-2 spike protein as sensor with little influence on the binding to hACE2 (Figure 1D) and minimal destruction of antibody-binding epitopes (Figure 4C); (ii) the acid-tolerant GFP-tag (mGam) enables fluorescent tracking and imaging analysis of dynamic viral entry events, even when it is internalized into acidic organelles (Figure S11B); (iii) the 293T-ACE2iRb3 cells with stably expressing ACE2-mRuby3 and H2B-iRFP670, allowing accurate membrane/nucleus definition and quantitative analysis at the single-cell level; (iiii) one-step, wash-free and fast detecting procedure (Figure 2D) provides robust settlement without biosafety concerns for high-throughput screening of neutralization antibodies and compound inhibitors.

Our data provided convincing evidence demonstrating the versatile applicability of the new system. The CSBT-determined entry-blocking potency was a better correlate of NAT against pseudotyping or the authentic SARS-CoV-2 virus than ELISA-binding activity in COVID19-convalescent human plasmas, immunized mouse sera and mAbs. The CSBT may serve as rapid proxy assessment to identify plasma source with therapeutic potential in clinic, and is a useful tool in evaluating vaccine efficacy and neutralizing mAb identification. In this study, 4 of 18 mAbs (36H6, 2B4, 3C8 and 12H8, Figure 4D) with the strongest CSBT blocking activities (IC50<10 nM) showed the most potent LVppNAT (IC90<3 nM). Notably, the 36H6 mAb presented superior neutralization activity against authentic SARS-CoV-2 (IC50=0.079 nM), which was comparable with recently described potent neutralizing mAbs^16, 31^, and thereby providing an excellent candidate for further development of therapeutic COVID-19 antibody ^32^. The 2B4 mAb has cross-neutralization activity for both SARS-CoV-1 and SARS-CoV-2 and showed binding activity with RaTG13-CoV, suggesting that it recognize a SARS-CoVs shared neutralizing-epitope. Further identification of such epitopes recognized by 2B4-like mAbs may facilitate the development of universal vaccines against SARS-CoV-like viruses ^33^.

Antibodies can inhibit viral infection via various mechanisms following the steps of viral cellular entry ^34, 35^. The CSBT and CRBT assay in combination with neutralization test provide a toolbox to distinguish the acting step of antibody neutralization (Figure 5). First, the mAbs (e.g. 36H6) with CRBT activities implying blocking capabilities on initial viral cell-attachment via hindering RBD-hACE2 interaction. Second, mAbs with CSBT but no CRBT activity (e.g. 53G2) may inhibit viral infection at post-attachment endocytic internalization. Third, some mAbs (e.g. 83H7) have neither CRBT nor CSBT effects and may also neutralize viruses via intracellular pathways when they enter cells by binding with viral spikes. It is possible that the intracellular antibody may block conformational changes and/or the requisite interaction between viral spike and host factor for viral-endolysosomal membrane fusion and/or viral genome release ^36^.

Furthermore, we developed STG-based high-content analysis system for studying compound-induced influences on SARS-CoV-2 endocytosis and intracellular trafficking (Figure 6). Our data revealed both CME and micropinocytosis involve in the STG entry process, whereas caveolae-mediated endocytosis plays little role. However, neither the CME nor micropinocytosis inhibitors completely blocked STG internalization and LVpp infection alone, suggesting that SARS-CoV-2 may use multiple mechanisms to gain entry into cells. These findings are consistent with previous studies regarding SARS-CoV-1 cell entry ^37^. The profound inhibitory effects of the acidification inhibitor Baf.A1 on both STG internalization and LVpp infection demonstrated that SARS-CoV-2 cell entry is pH-sensitive. In agreement with findings from a recent study, the inhibitors against TPC2 and PIKfyve strongly disturbed cell entry of STG and pseudotyped virus ^20^, showing potential drug targets for SARS-CoV-2 infection.

In summary, we developed a versatile tool for live-cell imaging studies of SARS-CoV-2 cell entry and provided a virus-free high-throughput assay to identify and characterize neutralizing antibodies and compound inhibitors. The new strategy can be adapted to develop visualization systems for studies cell entry of different viruses.

## Acknowledgments

This study was supported by National Natural Science Foundation of China (81993149041 for N.X.; 81902057 for Y.Z.; 81871316 and U1905205 for Q.Y.), the National Science and Technology Major Project of Infectious Diseases (No. 2017ZX10304402-002-003 for T.C. and No. 2017ZX10202203-009 for Q.Y.), the National Science and Technology Major Projects for Major New Drugs Innovation and Development (No. 2018ZX09711003-005-003 for T.C.), the Science and Technology Major Project of Fujian (2020YZ014001), the Science and Technology Major Project of Xiamen (3502Z2020YJ01) and the Guangdong Basic and Applied Basic Research Foundation (2020A1515010368 for C.S.). We thank Meng Lai, Wenyuan Liu and Hai Lin from PerkinElmer for technical assistance in imaging analysis and equipment maintenance.

## Author contributions

Y.Z., Q.Y., T.C., and N.X. conceptualized the project and designed the experiments. S.W., M.W., Z.L., S.N., and L.Z. expressed and purified the proteins. YTW., J.X., and Y.S. performed mouse immunization and developed monoclonal antibodies. Y.Z., J.W., J.Y., and L.Y., generated stable cell lines and performed cell imaging studies. Y.Z., H.H., J.M., and B.F. constructed plasmids for protein expression and cell experiments. Y.Z., H.H., J.W., J.Y., and H.X. performed the pseudovirus neutralization assays. C.S. performed the neutralization tests against the authentic virus in the P3 laboratory. S.W., M.W., and L.Z. performed immunoassays and SPR analyses. Q.Z. performed the cryo-EM reconstruction. Z.L. collected convalescent patients’ blood samples. H.Y. help to prepare the figures. Y.Z., L.Y., T.C., and Q.Y. integrated the data and wrote the manuscript. T.Y., L.L. J.H., S.G., Y.C., T.Z. and J.Z. provided critical revision of the manuscript for important intellectual content. Q.Y., T.C., and N.X. approved the final version of the manuscript.

## Methods

### Plasmas of convalescent COVID-19 patients

Plasma samples of a total of 32 convalescent COVID-19 patients were involved in this study. All of these patients were confirmed COVID-19 cases, and their samples were collected after they were discharged from the first hospital of Xiamen University. The study was approved by the institutional review board of the School of Public Health in accordance with the Declaration of Helsinki, and written informed consent was obtained. The characteristics of the patients and their samples were presented in Table S1.

### Cell lines

The cell lines of 293T, H1299, H1299-ACE2hR, 293T-ACE2hR and 293T-ACE2iRb3 were Dulbecco’s modified Eagle medium (Sigma, D6429) supplemented with 10% fetal bovine serum (Thermo Scientific, 10099-141), 0.1 mM non-essential amino acids (Thermo Scientific,1140-050), and were incubated at 37C° and 5% CO2 in a humidified incubator. To ensure the stable expression of transfected constructs in cells, the culture medium was supplemented with blasticidin (10μg/mL) for H1299-ACE2hR and 293T-ACE2hR, and was supplemented with puromycin (1μg/mL) for 293T-ACE2iRb3, respectively. The ExpiCHO-S cells were cultured with ExpiCHO™ Expression Medium (Thermo Scientific) in stackable CO2 incubator shaker.

### Mammalian cell expression vectors and lentiviral vectors

For mammalian cell expression, two modified PiggyBac (PB) transposon vectors (MIHIPsMie and EIRBsMie) were constructed based on PB-CMV-MCS-EF1α-RedPuro (System Biosciences, PB514B2). The fragment of CMV-MCS-EF1α-RedPuro on this vector was removed by SfiI/ApaI digestion. The DNA fragment of hCMVmie-MCS-IRES-H2BiRFP670-P2A-Puro-BGH and hCMVmie-MCS-IRES-H2BmRuby3-P2A-BsR-BGH were synthesized (Generalbiol, Anhui, China) and were ligated into the parental PB vector to generate the MIHIPsMie vector and EIRBsMie vector, respectively. The hCMVmie is an optimized CMV promoter with synthetic intron and is derived from pEE12.4 vector (Lonza). The iRFP670 is a near-infrared fluorescent protein with the excitation/emission maxima at 643 nm/670 nm ^38^. The mRuby3 is an improved red fluorescent protein with the excitation/emission maxima at 558 nm/592 nm ^39^.

The codon-optimized RBD gene of SARS-CoV-2 (referring to MN908947.3) was obtained by primer-annealing, following a PCR reaction for introductions of an N-terminal B2M leader sequence and a C-terminal polyhistidine sequence. Human codon-optimized DNA encoding the fluorescent proteins of Gamillus, mNeonGreen, and the full-length encoding genes of SARS-CoV-2 (GenBank: MN908947.3) and RaTG13-CoV (GISAID: EPI_ISL_402131) were synthesized (Generalbiol, Anhui, China). The encoding genes of spike proteins of SARS-CoV-1, MERS-CoV, and HKU1-CoV were purchased from Sino Biological Inc. The expression vectors for SARS-CoV1-RBG, RaTG13-RBG, HKU1-RBG, MERS-RBG, SARS-CoV2-RBN, SARS-CoV2-RBD, SARS-CoV2-STG, SARS-CoV2-STN, SARS-CoV2-ST, and SARS-CoV2-SMG were constructed as the frame structure described in Figure 1A and cloned into the EIRBsMie vector, via using NEBuilder HiFi DNA Assembly Master Mix (New England Biolabs).

For lentiviral vectors, the pLVEF1αIHRB-ACE2hR and pLVEF1αmNG vectors were constructed on the pLV-EF1α-MCS-IRES-Bsd vector (Youbio, VT8179). The ACE2 cDNA fragment (Sino Biological, HG10108-ACG) linking with an IRES-H2BmRuby3-P2A-BsR DNA fragment (synthesized by Generalbiol, Anhui, China) was cloned in-frame into the XbaI/SalI sites of pLV-EF1α-MCS-IRES-Bsd to obtain the pLVEF1αIHRB-ACE2hR vector.

### Recombinant proteins

Recombinant expressions of proteins involved in this study were performed by using the ExpiCHO™ expression system (Thermo Scientific, A29133). Briefly, plasmids encoding targeted proteins were transiently transfected into ExpiCHO-S cells by using ExpiFectamine™ CHO transfection kit (Thermo Scientific, A29129). Transfected cells were cultured in stackable CO2 incubator shaker (Kühner AG, SMX1503C). Cultures were harvested 5-7 days after transfection, and the cell-free supernatants were obtained by centrifugation and filtration with a 0.22 μm filter. Subsequently, the proteins in supernatants were captured by Ni Sepharose Excel resin, followed a wash with PBS buffer (20 mM PB7.4, 150 mM NaCl) containing 30 mM imidazole. Purified proteins were collected via a further elution with PBS buffer containing 250 mM imidazole, and were exchanged into the imidazole-free PBS buffer.

### Characterization of recombinant proteins by PAGE and SEC

Purified proteins were submitted to SDS-PAGE using SurePAGE (Genscript). Fluorescence detection in gel electrophoresis (Figure S1A) was performed using 1% agarose gel in 1x TAE buffer. Fluorescent gel image was acquired in FUSION FX7 Spectra multispectral imaging system (VILBER). The size exclusion liquid chromatography (SEC) for the SARS-CoV2-ST, SARS-CoV2-STG, and SARS-CoV2-STN proteins were performed using a high-performance liquid chromatography system (Waters Acquity UPLC) on a TSKgel G3000PWXL column. A gel filtration calibration HMW kit (GE health) was used for molecular weight calculation.

### Cryo-EM sample preparation, data collection, and processing

Aliquots (3 μL) of purified proteins of SARS-CoV2-ST or SARS-CoV2-STN were loaded onto glow-discharged (60 s at 20 mA) holey carbon Quantifoil grids (R2/1, 200 mesh, Quantifoil Micro Tools) using a Vitrobot Mark IV (ThermoFisher Scientific) at 100% humidity and 4°C. Data were acquired using the EPU software to control an FEI Tecnai F30 transmission electron microscope (Thermo Scientific) operated at 300 kV and equipped with a ThermoFisher Falcon-3 direct electron detector. Images were recorded in the 39-frames movie mode at a nominal magnification of 93,000X with a pixel size of 1.12 Å. The total electron dose was set to 30 e^−^Å^−2^ and the exposure time was 1s. Micrographs were collected with a defocus range comprised between 1.0 and 3.5 μm. Movie frame alignment and contrast transfer function estimation of each aligned micrograph were carried out with the programs Motioncor ^40^ and GCTF ^41^. Particle picking, two rounds of reference-free 2D classification and final 3D reconstruction were performed by the programs cryoSPARC v2 ^42^. Density-map-based visualization and segmentation were performed with Chimera ^43^.

### Generation and production of antibodies against SARS-CoV-2 S

Balb/c mice were intraperitoneal immunized with 5 μg of SARS-CoV2-RBD (expression in this study, n=5), SARS-CoV2-S1 (Sino Biological, 40591-V08H, n=3) and SARS-CoV2-S2 (Sino Biological, 40590-V08B, n=3), respectively. The proteins were emulsified in aluminum adjuvant for immunization. Triple immunizations were performed at week 0, 2, and 4. Two-week after immunization completion, mouse serum samples were collected for analyses as shown in Figure S4.

The mAbs against RBD of SARS-CoV-2 were raised in Balb/c mice using an injection 200 μg of SARS-CoV2-RBD protein emulsified in Freund’s complete adjuvant, followed by an intravenous booster injection of 200 μg of protein emulsified in Freund’s incomplete adjuvant at 2-week later, as previously described. The resulting hybridomas were screened for the secretion of RBD-specific mAbs using an indirect ELISA. The reactive cell clones were cultured in 75-cm^2^ flasks. Monoclonal cells that produced mAbs were obtained by limiting dilution at least three times. In this study, a total of 18 mAb-producing hybridomas were finally obtained. The mAbs were produced and purified as previously described ^44^.

### Enzyme-linked immunosorbent assay and western blots

The titers of TAb, IgG, and IgM against SARS-CoV-2 of human blood samples were detected by commercial enzyme-linked immunosorbent assay (ELISA) kits provided by Beijing Wantai Biological Pharmacy Enterprise Co.,Ltd. The measurements were performed according to the manufacturer’s instructions. The TAb-ELISA kit is based on recombinant viral antigen using a double-sandwich reaction form. The IgG kit is an indirect ELISA assay, and the IgM kit is based on the μ-chain capture method. All three assays used recombinant SARS-CoV-2 RBD antigens. The samples were initially tested undiluted, and the positive samples with the signal to a cutoff ratio (S/CO) >=10 were further diluted (1:10, 1:100, 1:1,000 and 1:10,000) by PBS buffer containing 20% newborn bovine serum (NBS) and tested again. The titers for TAb, IgG, and IgM antibody were calculated via S/CO multiplied by the maximum dilution factors.

To determine the ELISA binding activities of mAb to immobilized SARS-CoV2-RBD and SARS-CoV2-ST (Figure S5A), ELISA plates were coated with viral proteins at 200 ng per well, and nonspecific binding was blocked with phosphate-buffered saline (PBS) that contained 10% NBS, 0.5% casein (Sigma) and 10% sucrose. A series of 3-fold series dilutions that ranged from 10,000 ng/mL to 0.056 ng/mL for each mAb were prepared. For the test, 100 μl of specimens were added to the reaction well and incubated for 60-min at 37°C, followed by washing and reaction with horseradish peroxidase (HRP)-conjugated anti-mouse pAb (Wantai, Beijing, China). After a further 30-min incubation, the plates were washed with PBST buffer (20 mM PB7.4, 150 mM NaCl and 0.05% Tween 20) five times. The TMB chromogen solution (100 μL per well) was then added to the wells, and the plates were further incubated for 15-min. Subsequently, the reaction was stopped by adding 50 μL of 2 M H_2_SO_4_, and the OD was measured at 450 nm against 630 nm (OD_450-630_) by a microplate reader.

Epitope binning assays for mAbs (Figure S5B) were based on cELISA experiments. In brief, 96-well microplates were coated with SARS-CoV2-RBD at 200 ng per well. Aliquots of competitor mAbs (50 μL,10 μg per well) and HRP-conjugated mAbs (50 μL,10 μg per well) were added to the wells. The sample-loaded microplate was incubated at 37 °C for 1-hour. Then the microplate was washed five times with PBST buffer following the TMB chromogen solution addition. After a 15-min incubation, 50 μL of 2 M H_2_SO_4_ was added to stop the reaction, and the OD_450-630_ was measured. The inhibition ratio (%) was quantitatively assessed by comparing OD_450-630_ obtained with HRP-mAb in the presence or absence of competitor mAbs. A reduction of >70% was considered as an effective inhibition. The mAb clusters were generated based on the inhibition data by using HemI software ^45^.

Commercial antibodies were used to detect intracellular ACE2 (Sino Biological, 10108-T56), TMPRSS2 (Abcam, ab92323), and GAPDH (Proteintech, HRP-60004) according to the manufacturer’s instructions. The blots were imaged using FUSION FX7 Spectra multispectral imaging system (VILBER).

### Affinity determination and competition experiments using SPR

For determinations of the binding affinities of SARS-CoV2-RBG and SARS-CoV2-STG to hACE2 (Figure 1D), rACE2 (mouse-Fc tagged, Sino Biological) proteins were immobilized to a protein A sensorchip a level of ~500 response units (RUs) using Biacore 8000 (GE Healthcare) and a running buffer of composed of 20mM PB7.4 with 300 mM NaCl. Serial dilutions of purified SARS-CoV2-RBG and SARS-CoV2-STG proteins were injected ranging in concentration from 200 to 3.13 nM. To measure the affinities of mAbs to RBD proteins (Figure S6 and S10), various mAbs were loaded onto a protein A sensorchip to a level of ~1000 RUs and a running buffer of 20mM PB7.4. Serial dilutions of proteins (SARS-CoV2-RBD, SARS-CoV1-RBG or RaTG13-RBG) were injected ranging in concentration from 200 to 0.19 nM. The response data were fit to a 1:1 binding model using Biacore™ Insight evaluation software (GE Healthcare). For Fab competition experiments (Figure S8), rACE2 protein was loaded onto a protein A sensorchip at 200 nM. Subsequently, the SARS-CoV2-RBD protein (200 nM) were loaded to bind with rACE2 in the presence of 2-fold serial dilutions of various Fabs in concentration from 800 nM to 0 nM.

### Neutralization assays against pseudotyped and authentic virus

The SARS-CoV-2 and SARS-CoV-1 LVpp productions and LVppNAT measurements for blood samples and antibodies were performed as previously described ^46^. For determinations of compound-mediated inhibition for SARS-CoV-2 LVpp infection, the plated H1299-ACE2hR cells were pretreated with serial dilutions of compounds for 1-hour and then were incubated with LVpp inoculum (0.5 TU/cell). The cells were further cultured for 36-hour in the presence of compounds. Then the fluorescent imaging analysis and IC50 calculations were based on the infection-inhibition ratio of serial dilutions and determined by the 4-parameter logistic (4PL) regression using GraphPad Prism v8.0. Neutralization activities of COVID-19-convalescent human plasmas and mAbs against authentic SARS-CoV-2 virus were detected as previously described ^25^. Briefly, 2-fold serial dilutions of plasma samples (from 1:10 to 1:10240) and mAbs (from 100 μg/mL to 0.763 ng/mL) were prepared and incubated with 100 times the tissue culture infective dose (TCID50) of the BetaCoV/Shenzhen/SZTH-003/2020 strain virus (GISAID access number: EPI_ISL_406594) at 37°C for 1-hour. The mixtures were then added to a monolayer of Vero cells (10^4^ cells per well, pre-washed twice with PBS) in a 96-well plate and incubated at 37°C. Microscopic examinations were performed for the cytopathic effect after 5-day incubation. The complete absence of cytopathic effect in an individual culture well was defined as protection. The ID50 (for plasma samples) or IC50 (for mAbs) were calculated using GraphPad Prism.

### Cell imaging assays

For direct visualizing the cellular binding and uptake of RBD or spike proteins, the 293T-ACE2iRb3 cells were seeded at 2×10^4^ cells per well in poly-D-lysine pretreated CellCarrier-96 Black plate. After 1-day culture, the fluorescent probes (ensure a final concentration of 25 nM for RBD based protein probes or 2.5 nM for ST based protein probes in culture medium) were added to the cell cultures. In experiments of Figure 2B, the cells were cultured at 37°C in CO2 incubator for 0, 6, 30, 60, and 120 -min, and were gently washed twice with PBS following a paraformaldehyde fixation. The images of Figure 2B were acquired on TCS SP8 STED confocal microscope (Leica Microsystems) using a 100x oil immersion objective. In experiments of Figure S3A-D, the cell culture plate (after probe loading, in live-cell and wash-free conditions) was placed in a pre-heated (37°C) Opera Phenix with 3% CO2. Multi-channel fluorescence (STG or RBG, Ex:488/Em:525; ACE2-mRuby3, Ex:561/Em:590; H2BiRFP670, Ex:640/Em670) cell images were acquired every 6-min (0 to120-min). In experiments of Figure 2D-F, cell images were acquired (Opera Phenix) at 1-hour after probe loading in wash-free and live-cell conditions.

For CSBT and CRBT assays, blood samples or mAbs were pre-made as 2-fold serial dilutions using DMEM containing 10% FBS. Aliquots (44 μL per well) of diluted samples and protein probes (11 μL per well) were mixed in a 96-well plate with U shaped bottom. Half of the culture medium (50 μL) of 293T-ACE2iRb3 cell plate was gently removed, and 50 μL of sample/probe mixtures were added to each well. Cell image acquisitions performed with Opera Phenix (green, red and near-infrared channels in confocal mode) using a 20x water immersion objective at 1-hour after probe incubation in wash-free and live-cell conditions.

In simultaneous tracking of STG and mAbs (Figure 5), the 293T-ACE2iRb3 cells in CellCarrier-96 Black plate were pre-stained with NucBlue before the incubations of mAbs and STG. Subsequently, aliquots (10 μL) of dylight633-labeled mAbs of 36H6, 53G2, 83H7, 8H6 and ctrAb (to achieve a final concentration of 20 nM) with 2.5 nM (a final concentration in culture medium) of STG probe (10 μL) or not, were added into to the wells, respectively. The plate was immediately placed in a pre-heated (37°C) Opera Phenix with 3% CO2. Time-serial four-channel (NucBlue: Ex:405/Em:450; STG or RBG, Ex:488/Em:525; ACE2-mRuby3, Ex:561/Em:590; H2BiRFP670, Ex:640/Em670) live-cell images were acquired at 10-min, 1-hour, 2-hour, 3-hour, 5-hour, 7-hour, 9-hour, 11-hour, and 13-hour using a 63x water immersion objective.

To visualize compound-induced influence on viral entry, 293T-ACE2iRb3 (Figure 6C-D, Figure S11A) or H1299-ACE2hR cells (Figure S11B) were pretreated with serial dilutions of compounds for 1-hour. Then the probes were added to the cell cultures for further incubations in the presence of compounds. Cell images shown in Figure 6C and Figure S11B were acquired on TCS SP8 STED confocal microscope using a 100x oil immersion objective. The data of Figure 6D and Figure S11A were derived from images acquired on Opera Phenix using 40x water immersion objective. For pictures of Figure 6C, the cells were gently washed twice with PBS at 5-hour post STG incubation, following a paraformaldehyde fixation before imaging. For experiments as shown in Figure S11B, the cells at 5-hour post STG incubation were stained with Lysoview633 (0.1 μL per well) for 10-min, then the cells were gently washed twice with PBS buffer and fixed with paraformaldehyde treatment before imaging. Cell images involved in Figure 6D and Figure S11A were acquired in wash-free and live-cell conditions, at different various time points as indicated in their legends.

### Quantitative image analyses

All quantitative image analyses were based on images acquired by Opera Phenix, following a schematic flow chart shown in Figure 2D. All image data were transfer to Columbus system (version 2.5.0, PerkinElmer Inc) for analysis. Multiparametric image analysis was performed as described in the following. The signals of the blue channel (NucBlue, only for Figure 5) or near-infrared channel (H2BiRFP, for other data) were used to detect the nucleus. As the ACE2 is a membrane protein, the signals of ACE2-mRuby3 (red channel) were used to determine the cell boundary. Then, the cells were further segment into the regions of membrane (outer border: 0%, inner border: 15%), cytoplasm (outer border: 20%, inner border: 45%), and nucleus (outer border: 55%, inner border: 100%). For CSBT and CRBT assays, the MFI of probe channel (Ex488/Em525) in the cytoplasmic region (cMFI). The MFI of ACE2-mRuby3 (Ex561/Em590) on the membrane were also calculated for inter-well normalization. The cMFI inhibition ratio (%) of the test sample was calculated using the following equation: [(cMFI_pc_-cMFI_tst_)/(cMFI_pc_-cMFI_blk_)]×100%. In this formula, the cMFI_pc_ is the cMFI value of probe-only well (as positive control), the cMFI_tst_ is the cMFI value of test well, and the cMFI_blk_ is the cMFI value of cell-only well. For each plate, five replicates of probe-only well and one cell-only well were included. The CSBT and CRBT activities of mAbs were expressed as IC50, and that of blood samples were expressed as ID50. The ID50/IC50 values were determined by 4PL regression GraphPad Prism v8.0. To determine the internalization characteristics, the parameters of IFR, IVNs, IVpMFI, and IVA were measured. Among these parameters, the IFR is the ratio of intensity in the cytoplasmic region and in the whole cell, the IVNs is the average numbers (per cell) of internalized fluorescent vesicles, the IVpMFI is the average peak MFI of internalized fluorescent vesicles, and the IVA is the average area of internalized fluorescent vesicles. The detailed algorithms for the above-mentioned imaging analyses using Columbus system are available from the corresponding authors on request.

### Statistical analysis

The unpaired t-test of variance was used to compare continuous variables. Linear regression models and Pearson correlation tests were used for correlation analyses. Two-way ANOVA tests were used to analysis the time-serial observations for independent variable. Differences were considered significant at a two-tailed p < 0.05. GraphPad Prism version 8.0.1 was used for all statistical calculations.

---

**Figure S1. SDS-PAGE analyses of various recombinant proteins involved in the study.** Related to Figure 1. (A) SDS-PAGE (left panel) and fluorescence naïve PAGE (right panel) of SARS-CoV2-RBG. Mr: molecular weight marker; lane 1: supernatants from transfected cells; lane 2: supernatants after flowing through Ni Sepharose Excel resin; lane 3: wash fraction with 30 mM imidazole; lane 4: reduced lane 3 sample; lane 5: elution with 250 mM imidazole; lane 6: reduced lane 5 sample. (B-K) SDS-PAGE analyses of SARS-CoV1-RBG (B), RaTG13-RBG (C), HKU1-RBG (D), MERS-RBG (E), SARS-CoV2-RBN (F), SARS-CoV2-RBD (G), SARS-CoV2-STG (H), SARS-CoV2-STN (I), SARS-CoV2-ST (J), and SARS-CoV2-SMG (K). Mr: molecular weight marker; lane 1: supernatants from transfected cells; lane 2: supernatants after flowing through Ni Sepharose Excel resin; lane 3: wash fraction with 30 mM imidazole; lane 4: elution with 250 mM imidazole. The bands corresponding to targeted protein are denoted with red arrows.

**Figure S2. Protein characterizations by SEC and Cryo-EM.** Related to Figure 1. (A) SEC chromatograms of 7 protein standards on a TSK-G3000 column. (B) A calibration curve based on data of (A) for calculation protein molecular weight. Cryo-EM reconstructions of CHO-derived SARS-CoV2-ST (C) and SARS-CoV2-STN (D) proteins. Ten representative 2D classification averages illustrating particles with prefusion orientations (upper panel). The 8.9-Å density map of SARS-CoV2-ST, and the ~22-Å density map of SARS-CoV2-STN were colored by protomers, respectively (lower panel).

**Figure S3. Live-cell imaging analyses in comparison of the probes of STG and RBG of SARS-CoV-2.** (A) Time-lapse live-cell images from a single observation on 293T-ACE2iRb3 cells incubated with SARS-CoV2-STG (upper panel) and SARS-CoV2-RBG (lower panel). Scale bar, 20 μm. The cell nucleus H2B-iRFP670 was pseudo-colored blue. Quantitative comparisons of the internalized fluorescence intensity ratios of mGam (B) and ACE2-mRuby3 (C) in between 293T-ACE2iRb3 cells incubated with SARS-CoV2-RBG and SARS-CoV2-STG during 6 to 126 minutes. Mock indicated untreated cells. Data were mean±SEM derived from time-lapse imaging of about 200 cells. (D) Split violin plots to compare the numbers of internalized mGam-active vesicles in between 293T-ACE2iRb3 cells incubated with STG and RBG at 6, 30, 60 and 120 minutes. (E) Quality measurements of the CSBT (left panel) and CRBT (right panel) assays. The Z’-factor was determined as described in the Methods section. Pos-CTR, positive control; Neg-CTR, negative control.

**Figure S4. Detections of antibody titers of immunized mouse sera by cell-based assays.** (A) LVppNAT on H1299-ACE2hR cells. (B) CSBT and (C) CRBT on 293T-ACE2iRb3 cells. (D) A summary of the titers (ID50) detected by LVppNAT, CSBT and CRBT. Serum samples for mice immunized with recombinant proteins of SARS-CoV2-S1 (mouse S1-1, S1-2, S1-3), SARS-CoV2-S2 (mouse S2-1, S2-2, S2-3) and SARS-CoV2-RBD (mouse RBD-1, RBD-2, RBD-2, RBD-3, RBD-4, RBD-5) were detected.

**Figure S5. ELISA analyses for mAbs.** (A) ELISA-binding activities of mAbs to immobilized SARS-CoV2-RBD (upper panel) and SARS-CoV2-ST (lower panel) proteins. Recombinant RBD and ST proteins of SARS-CoV-2 were coated on the ELISA plates at 200 ng/well. Different mAbs were tested at a 3-fold serial dilutions that began at 10,000 ng/mL. (B) Epitope binning assays for mAbs. A heatmap representation of a cross-competition ELISA with 18 mAbs developed in this study. The mAbs listed on the horizontal axis were conjugated with HRP and were used to react with the RBD-coated microplate. The mAbs listed on the vertical axis were the competitor mAb. A reduction of >70% of ELISA OD values of RBD-mAb-HRP capture in the presence of competitor mAb was considered as an effective inhibition. The mAb clusters were generated based on the inhibition data by using HemI software.

**Figure S6. SPR sensorgrams showing the binding kinetics for SARS-CoV2-RBD and immobilized mAbs.** Related to Figure 4C and Table S4. Colored lines represented a global fit of the data at known concentrations using a 1:1 binding model.

**Figure S7. Titrations of mAbs in CRBT, LVppNAT and authentic SARS-CoV-2 neutralization assays.** (A) Inhibition potencies of mAb, which have typical dose-dependent inhibitory effects, in CRBT assay. A broken line indicates the demarcation line of 50% inhibition. (B) Dose-response curves of mAbs with enhancement potential in CRBT assay. The upper broken line indicates the demarcation line of 50% inhibition, whereas the lower broken line indicates the demarcation line of 50% enhancement. (C) Neutralization potencies of the mAbs against SARS-CoV-2 measured by using LVppNAT. (D) Neutralization tests of the mAbs of 36H6, 2B4, 34B4, 83H7, 8H6, and CR3022 against SARS-CoV-1 LVpp. (E) Neutralization potencies of the 36H6 and 2B4 mAbs against authentic SARS-CoV-2 virus. ctrAb, control mAb.

**Figure S8. Biacore analyses for the influence of Fabs derived from various mAbs on the interaction between ACE2 and SARS-CoV2-RBD.** SPR sensorgrams showing the binding of SARS-CoV2-RBD (200 nM) to with immobilized human ACE2 (200 nM) in the presence of various Fabs at different concentrations (800, 400, 200, 100, 50, 25, 12.5, 6.25, and 3.13 nM of Fab were tested).

**Figure S9. Correlations between the LVppNAT and the activities measured by CSBT, CRBT, and ELISAs for mAbs.** The relationship between CSBT-IC50 and LVppNAT-IC50 (A). Correlations between CRBT-IC50 and LVppNAT-IC90 (B) or LVppNAT-IC50 (C). Correlations between LVppNAT-IC90 and RBD-binding affinity (D), RBD-ELISA binding activities (E), ST-ELISA binding activities of mAb (F), respectively.

**Figure S10. SPR sensorgrams showing the binding kinetics for 6 representative mAbs to SARS-CoV1-RBG (A) and RaTG13-RBG (B).** Related to Figure 4C and Table S4. The mAbs of 36H6, 2B4, 34B4, 5F3, 18C5, and 8H6 were tested at 2-fold serial dilutions (200, 100, 50, 25, 12.5, 6.25, 3.13, 1.56, 0.78, 0.39, and 0.19 nM).

**Figure S11. Characterization of compound-induced influence on STG internalization on ACE2-expressing cells.** (A) Comparisons of cytoplasmic STG intensity (cMFI) of 293T-ACE2iRb3 cells treated by various compounds. Related to Figure 6. Cell imaging was performed 1-hour post STG incubation and the cMFI were calculated following approach as described in Figure 2D. (B) Colocalizations of internalized STG vesicles and lysosomes in H1299-ACE2hR cells treated by various compounds. STG exhibited green fluorescent signal, the lysosome was stained with Lysoview633 showing red fluorescent signal. Fluorescent images were obtained at 5-hour post STG incubation. Both for (A) and (B), the cells were pretreated with compounds at their maximal non-cytotoxic concentrations for 1-hour before STG probe loading.

**Figure S12. Compound-induced changes of on STG-internalization related characteristics correlate with the inhibitory effects against SARS-CoV-2 LVpp infection.** Related to Figure 6. Correlations between the characteristic parameters of STG-IVNs (left panel), STG-IVA (middle panel), STG-IFR (right panel) at 1-hour (upper panel) or 5-hour (lower panel) (data were derived from Figure 6D), and the relative SARS-CoV-2 LVpp infection efficiency (%, data were derived from Figure 6B).

