## Supplemental Figures and Tables for "Virus-free and live-cell visualizing SARS-CoV-2 cell entry for studies of neutralizing antibodies and compound inhibitors"

Figure S1

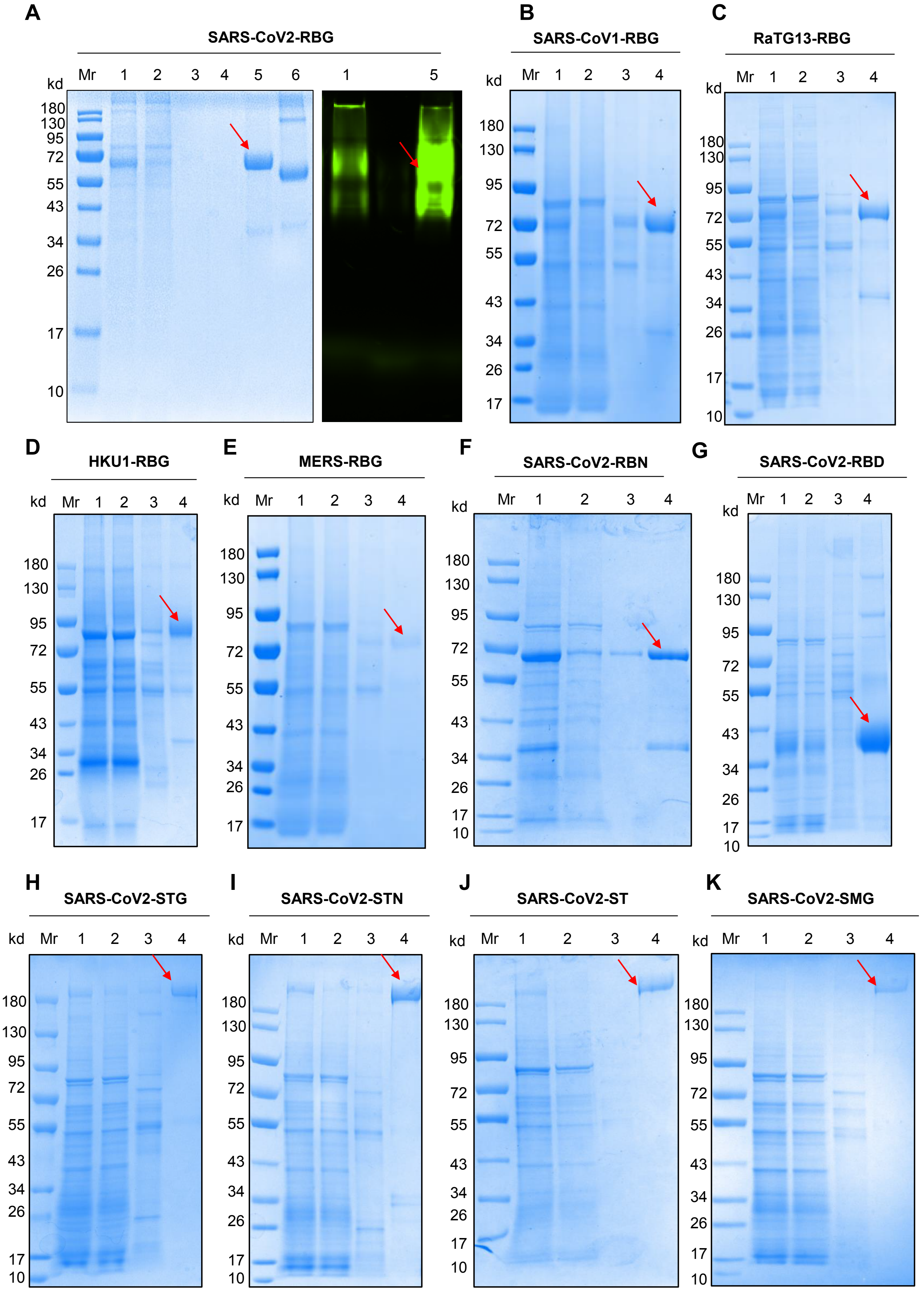
A
SARS-CoV2-RBG
kd
Mr 1 2 3 4 5 6 1 5
180
130
95
72
55
43
34
26
17
10
B
SARS-CoV1-RBG
C
RaTG13-RBG
D
HKU1-RBG
E
MERS-RBG
F
SARS-CoV2-RBN
G
SARS-CoV2-RBD
H
SARS-CoV2-STG
I
SARS-CoV2-STN
J
SARS-CoV2-ST
K
SARS-CoV2-SMG

#### Figure S2

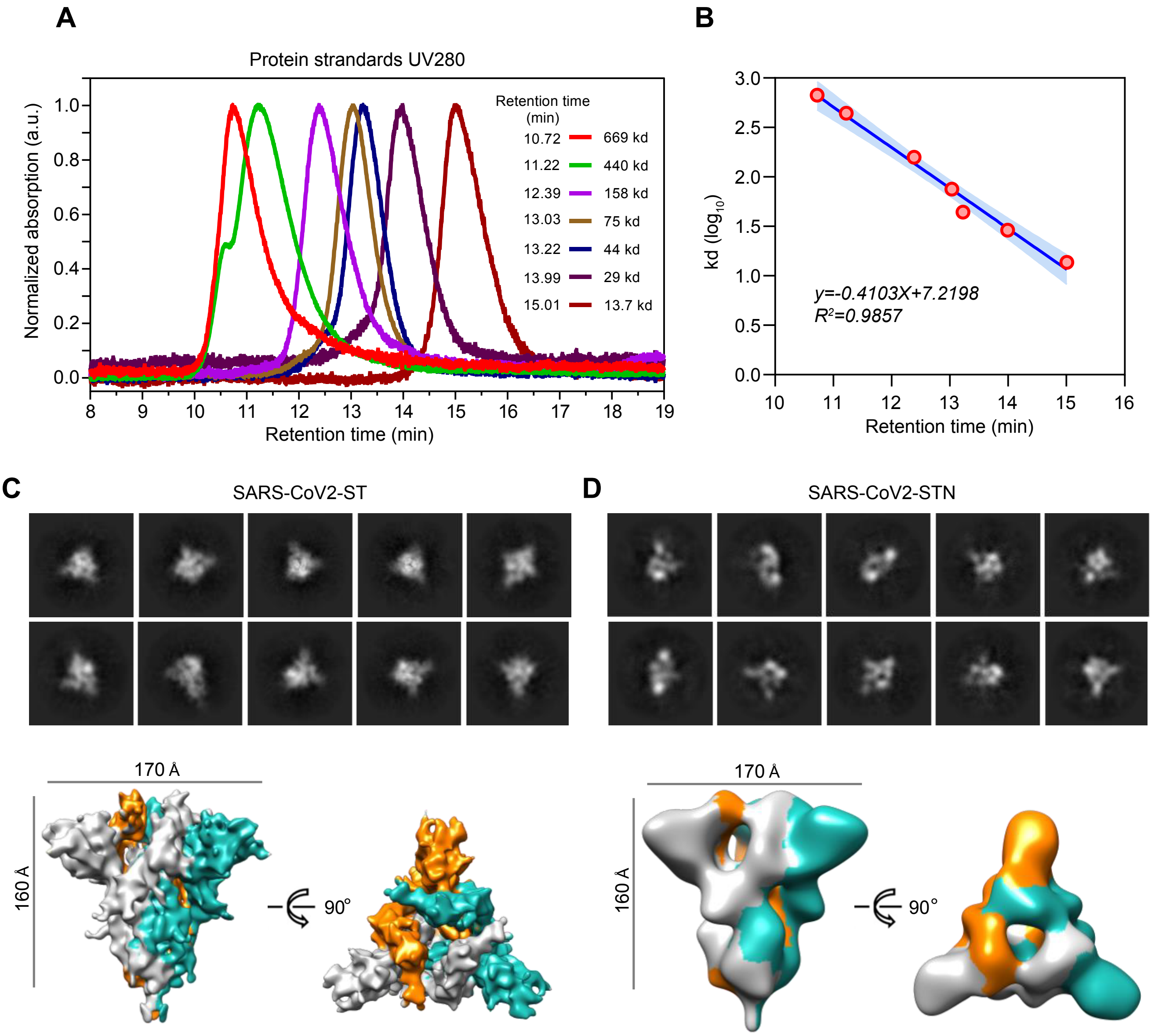

A
Protein strandards UV280
Normalized absorption (a.u.)
1.0
0.8
0.6
0.4
0.2
0.0
Retention time (min)
Retention time (min)
10.72 669 kd
11.22 440 kd
12.39 158 kd
13.03 75 kd
13.22 44 kd
13.99 29 kd
15.01 13.7 kd
B
kd ($log_{10}$)
3.0
2.5
2.0
1.5
1.0
0.5
0.0
10
11
12
13
14
15
16
Retention time (min)
y=-0.4103X+7.2198
$R^2$=0.9857
C
SARS-CoV2-ST
170 Å
160 Å
90°
D
SARS-CoV2-STN
170 Å
160 Å
90°

### Figure S3

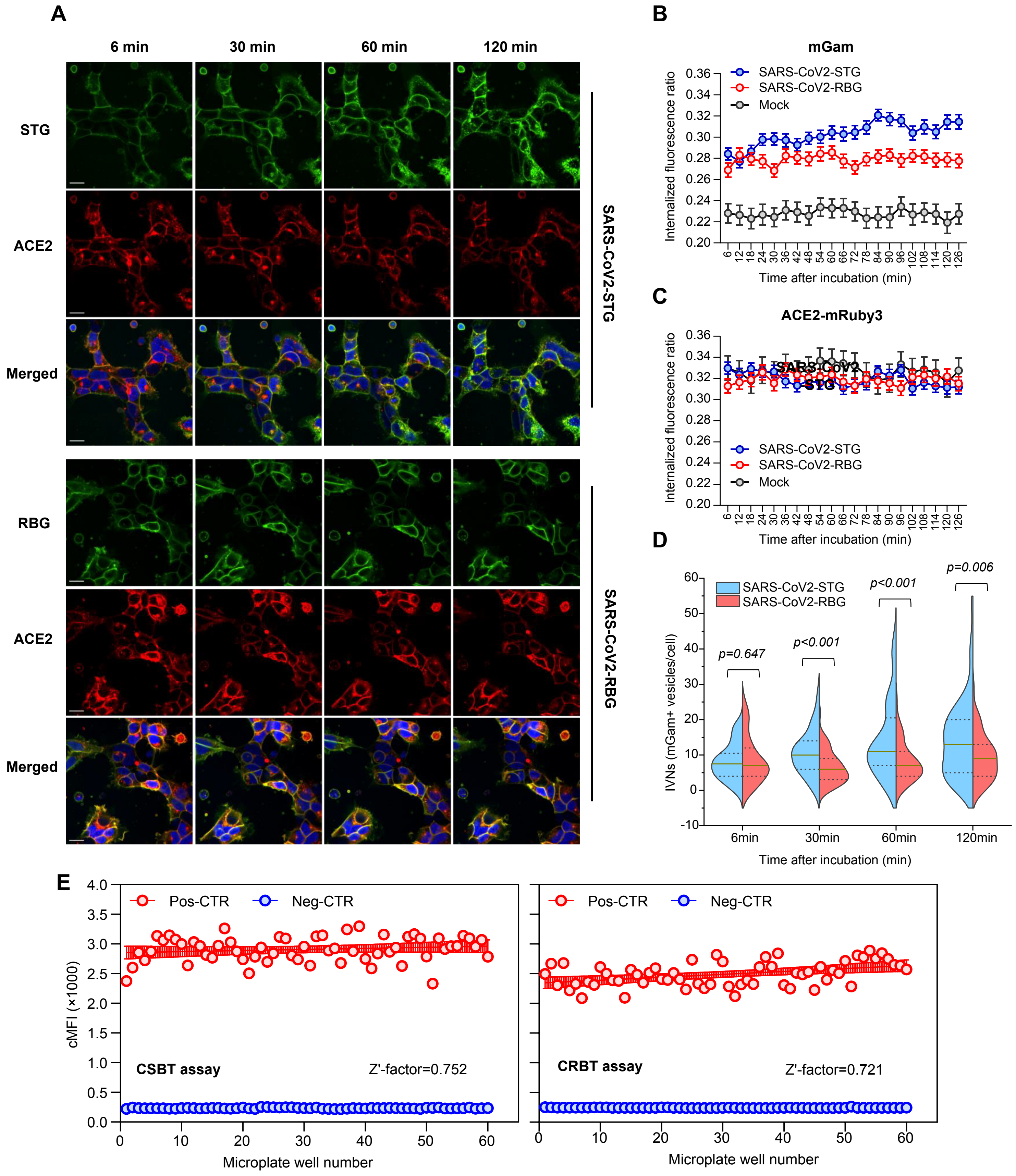

A
6 min
30 min
60 min
120 min
STG
ACE2
Merged
SARS-CoV2-STG
RBG
ACE2
Merged
SARS-CoV2-RBG
B
mGam
SARS-CoV2-STG
SARS-CoV2-RBG
Mock
Internalized fluorescence ratio
Time after incubation (min)
C
ACE2-mRuby3
SARS-CoV2-STG
SARS-CoV2-RBG
Mock
Internalized fluorescence ratio
Time after incubation (min)
D
SARS-CoV2-STG
SARS-CoV2-RBG
p=0.647
p<0.001
p<0.001
p=0.006
IVNs (mGam+ vesicles/cell)
6min
30min
60min
120min
Time after incubation (min)
E
Pos-CTR
Neg-CTR
cMFI (×1000)
CSBT assay
Z'-factor=0.752
Microplate well number
Pos-CTR
Neg-CTR
CRBT assay
Z'-factor=0.721
Microplate well number

### Figure S4

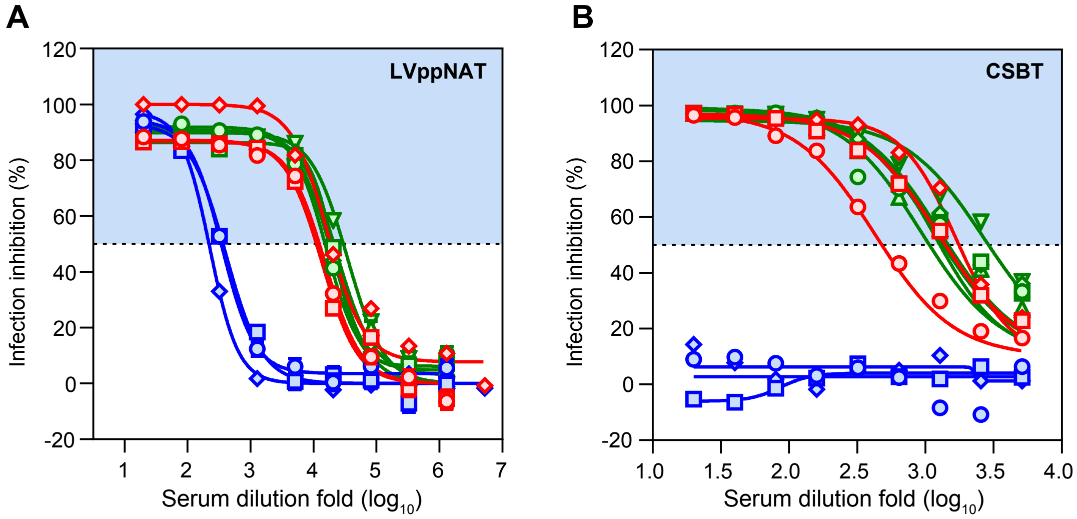

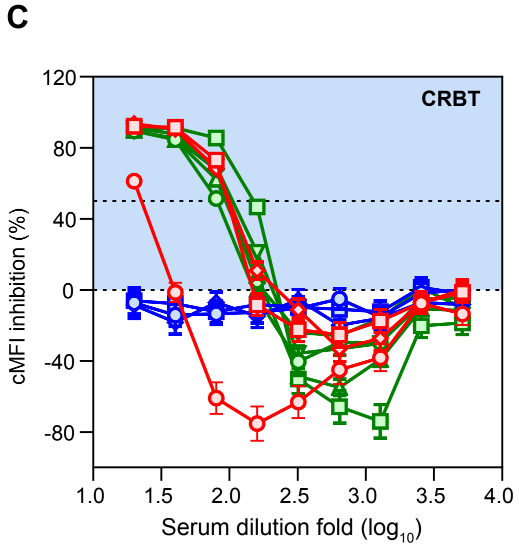

**D**

| Mouse ID | LVppNAT (ID50) | CSBT (ID50) | CRBT (ID50) |
|---|---|---|---|
| S1-1 | *13,563* | *427* | *23* |
| S1-2 | *15,164* | *1,259* | *84* |
| S1-3 | *16,843* | *1,663* | *105* |
| S2-1 | *351* | *<20* | *<20* |
| S2-2 | *403* | *<20* | *<20* |
| S2-3 | *226* | *<20* | *<20* |
| RBD-1 | *16,024* | *932* | *84* |
| RBD-2 | *21,458* | *1,186* | *170* |
| RBD-3 | *20,413* | *1,165* | *98* |
| RBD-4 | *23,327* | *1,285* | *100* |
| RBD-5 | *34,663* | *2,699* | *116* |

### Figure S5

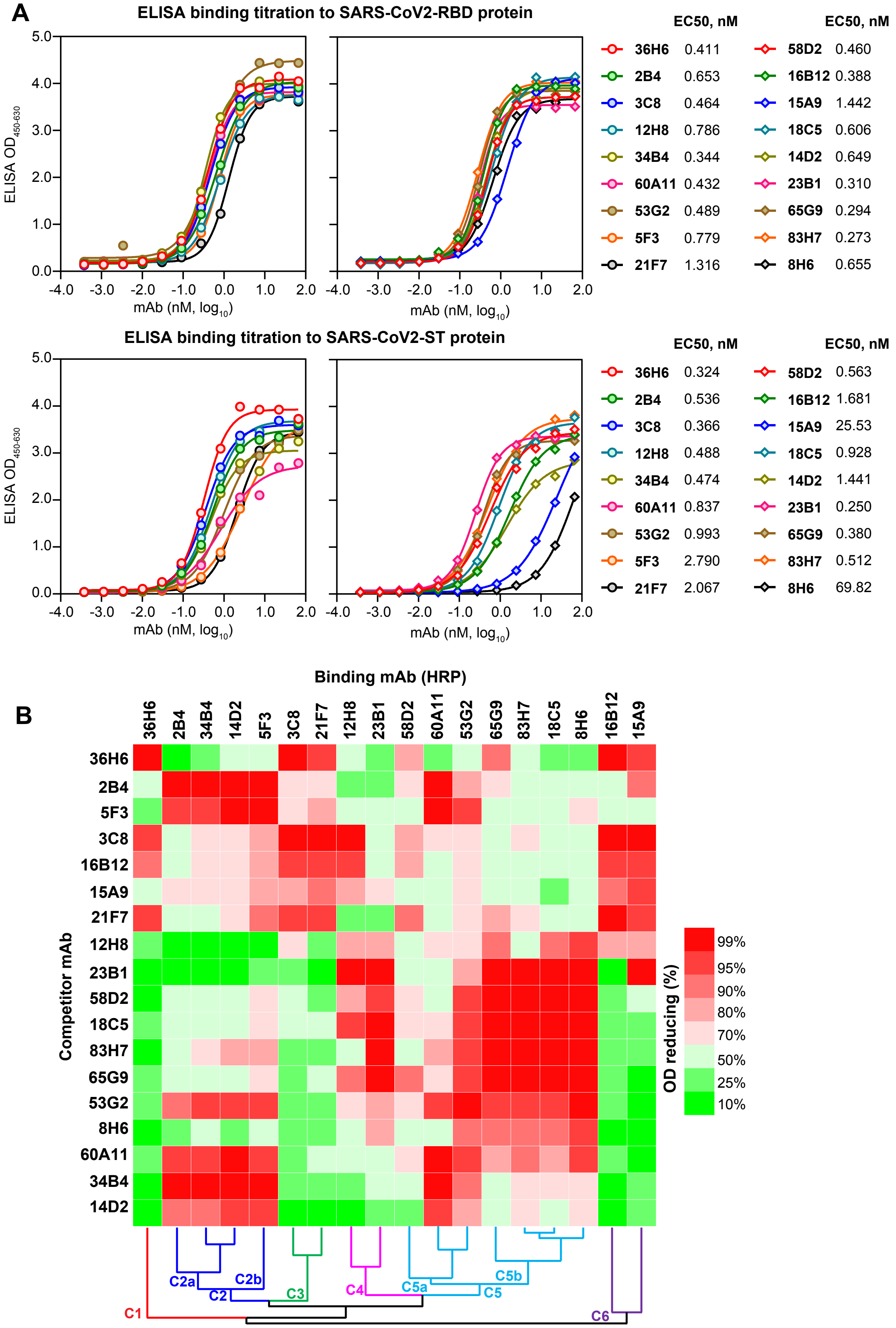

A
ELISA binding titration to SARS-CoV2-RBD protein
ELISA OD450-630
mAb (nM, log10)
EC50, nM
36H6 0.411
2B4 0.653
3C8 0.464
12H8 0.786
34B4 0.344
60A11 0.432
53G2 0.489
5F3 0.779
21F7 1.316
58D2 0.460
16B12 0.388
15A9 1.442
18C5 0.606
14D2 0.649
23B1 0.310
65G9 0.294
83H7 0.273
8H6 0.655
ELISA binding titration to SARS-CoV2-ST protein
36H6 0.324
2B4 0.536
3C8 0.366
12H8 0.488
34B4 0.474
60A11 0.837
53G2 0.993
5F3 2.790
21F7 2.067
58D2 0.563
16B12 1.681
15A9 25.53
18C5 0.928
14D2 1.441
23B1 0.250
65G9 0.380
83H7 0.512
8H6 69.82
B
Binding mAb (HRP)
36H6 2B4 34B4 14D2 5F3 3C8 21F7 12H8 23B1 58D2 60A11 53G2 65G9 83H7 18C5 8H6 16B12 15A9
Competitor mAb
36H6 2B4 5F3 3C8 16B12 15A9 21F7 12H8 23B1 58D2 18C5 83H7 65G9 53G2 8H6 60A11 34B4 14D2
OD reducing (%)
99% 95% 90% 80% 70% 50% 25% 10%
C1 C2 C2a C2b C3 C4 C5 C5a C5b C6

### Figure S6

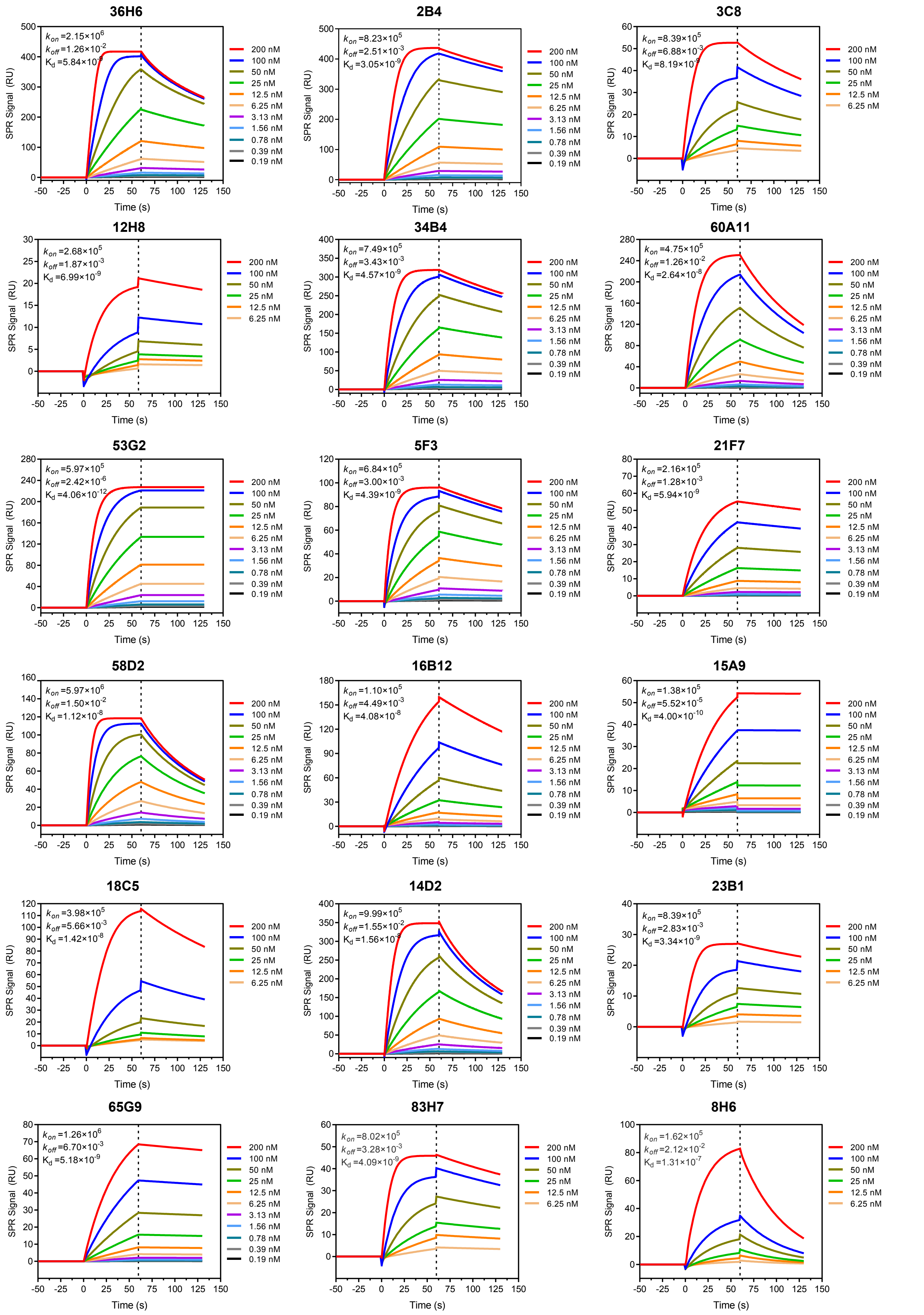

36H6
$k_{on}$ =2.15×10^6
$k_{off}$ =1.26×10^-2
$K_d$ =5.84×10^-9
2B4
$k_{on}$ =8.23×10^5
$k_{off}$ =2.51×10^-3
$K_d$ =3.05×10^-9
3C8
$k_{on}$ =8.39×10^5
$k_{off}$ =6.88×10^-3
$K_d$ =8.19×10^-9
12H8
$k_{on}$ =2.68×10^5
$k_{off}$ =1.87×10^-3
$K_d$ =6.99×10^-9
34B4
$k_{on}$ =7.49×10^5
$k_{off}$ =3.43×10^-3
$K_d$ =4.57×10^-9
60A11
$k_{on}$ =4.75×10^5
$k_{off}$ =1.26×10^-2
$K_d$ =2.64×10^-8
53G2
$k_{on}$ =5.97×10^5
$k_{off}$ =2.42×10^-6
$K_d$ =4.06×10^-12
5F3
$k_{on}$ =6.84×10^5
$k_{off}$ =3.00×10^-3
$K_d$ =4.39×10^-9
21F7
$k_{on}$ =2.16×10^5
$k_{off}$ =1.28×10^-3
$K_d$ =5.94×10^-9
58D2
$k_{on}$ =5.97×10^6
$k_{off}$ =1.50×10^-2
$K_d$ =1.12×10^-8
16B12
$k_{on}$ =1.10×10^5
$k_{off}$ =4.49×10^-3
$K_d$ =4.08×10^-8
15A9
$k_{on}$ =1.38×10^5
$k_{off}$ =5.52×10^-5
$K_d$ =4.00×10^-10
18C5
$k_{on}$ =3.98×10^5
$k_{off}$ =5.66×10^-3
$K_d$ =1.42×10^-8
14D2
$k_{on}$ =9.99×10^5
$k_{off}$ =1.55×10^-2
$K_d$ =1.56×10^-8
23B1
$k_{on}$ =8.39×10^5
$k_{off}$ =2.83×10^-3
$K_d$ =3.34×10^-9
65G9
$k_{on}$ =1.26×10^6
$k_{off}$ =6.70×10^-3
$K_d$ =5.18×10^-9
83H7
$k_{on}$ =8.02×10^5
$k_{off}$ =3.28×10^-3
$K_d$ =4.09×10^-9
8H6
$k_{on}$ =1.62×10^5
$k_{off}$ =2.12×10^-2
$K_d$ =1.31×10^-7
SPR Signal (RU)
Time (s)
200 nM
100 nM
50 nM
25 nM
12.5 nM
6.25 nM
3.13 nM
1.56 nM
0.78 nM
0.39 nM
0.19 nM

**Figure S7**

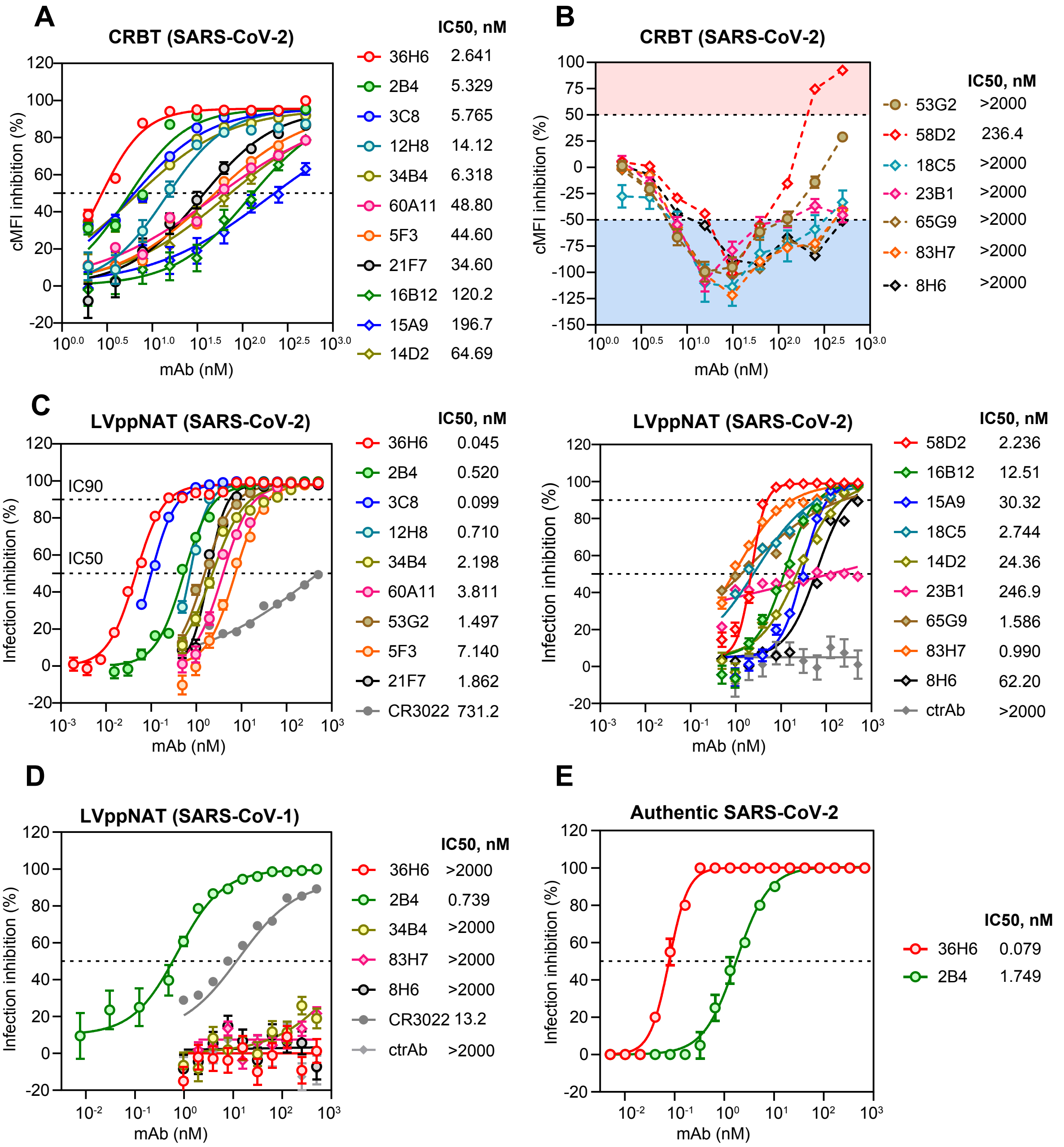
A
CRBT (SARS-CoV-2)
cMFI inhibition (%)
mAb (nM)
IC50, nM
36H6 2.641
2B4 5.329
3C8 5.765
12H8 14.12
34B4 6.318
60A11 48.80
5F3 44.60
21F7 34.60
16B12 120.2
15A9 196.7
14D2 64.69
B
CRBT (SARS-CoV-2)
cMFI inhibition (%)
mAb (nM)
IC50, nM
53G2 >2000
58D2 236.4
18C5 >2000
23B1 >2000
65G9 >2000
83H7 >2000
8H6 >2000
C
LVppNAT (SARS-CoV-2)
Infection inhibition (%)
IC90
IC50
mAb (nM)
IC50, nM
36H6 0.045
2B4 0.520
3C8 0.099
12H8 0.710
34B4 2.198
60A11 3.811
53G2 1.497
5F3 7.140
21F7 1.862
CR3022 731.2
LVppNAT (SARS-CoV-2)
Infection inhibition (%)
mAb (nM)
IC50, nM
58D2 2.236
16B12 12.51
15A9 30.32
18C5 2.744
14D2 24.36
23B1 246.9
65G9 1.586
83H7 0.990
8H6 62.20
ctrAb >2000
D
LVppNAT (SARS-CoV-1)
Infection inhibition (%)
mAb (nM)
IC50, nM
36H6 >2000
2B4 0.739
34B4 >2000
83H7 >2000
8H6 >2000
CR3022 13.2
ctrAb >2000
E
Authentic SARS-CoV-2
Infection inhibition (%)
mAb (nM)
IC50, nM
36H6 0.079
2B4 1.749

### Figure S8

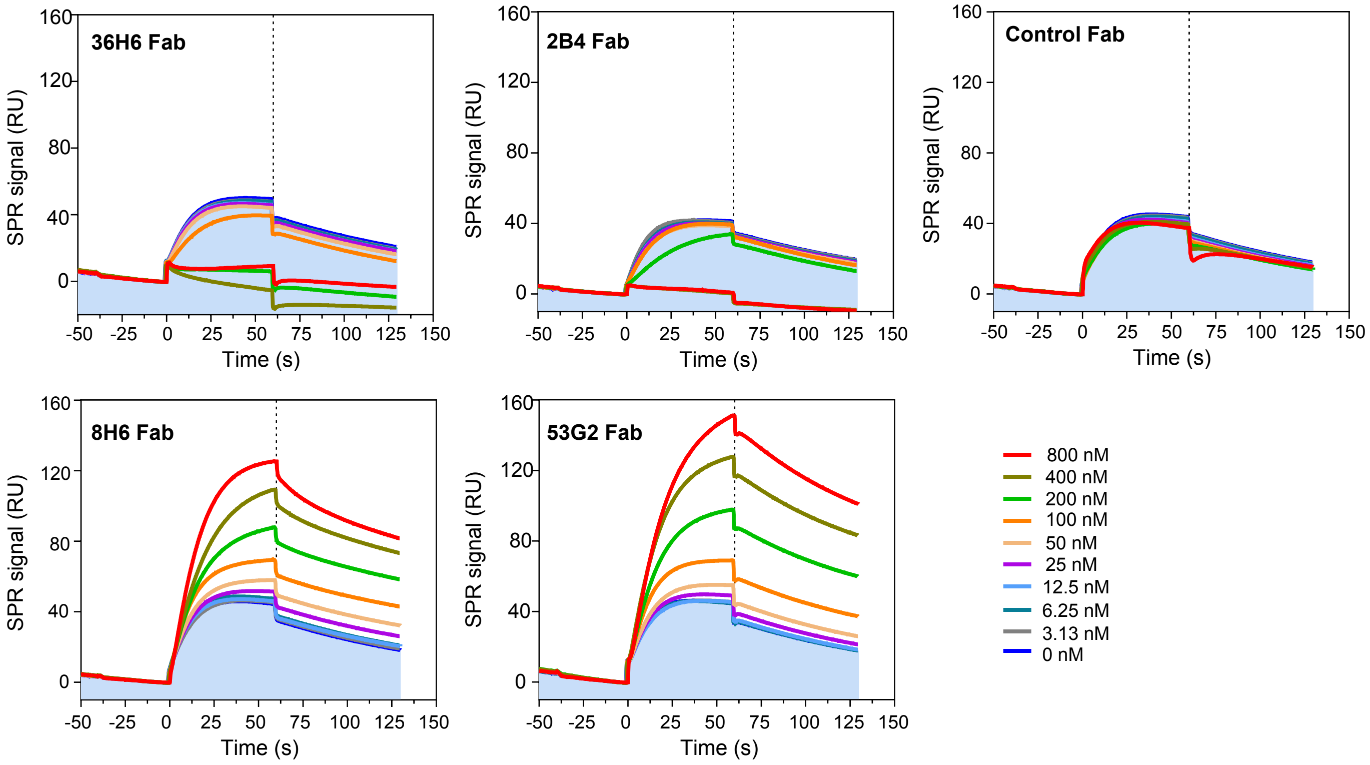

36H6 Fab
2B4 Fab
Control Fab
8H6 Fab
53G2 Fab
SPR signal (RU)
Time (s)
160
120
80
40
0
-50
-25
0
25
50
75
100
125
150
800 nM
400 nM
200 nM
100 nM
50 nM
25 nM
12.5 nM
6.25 nM
3.13 nM
0 nM

#### Figure S9

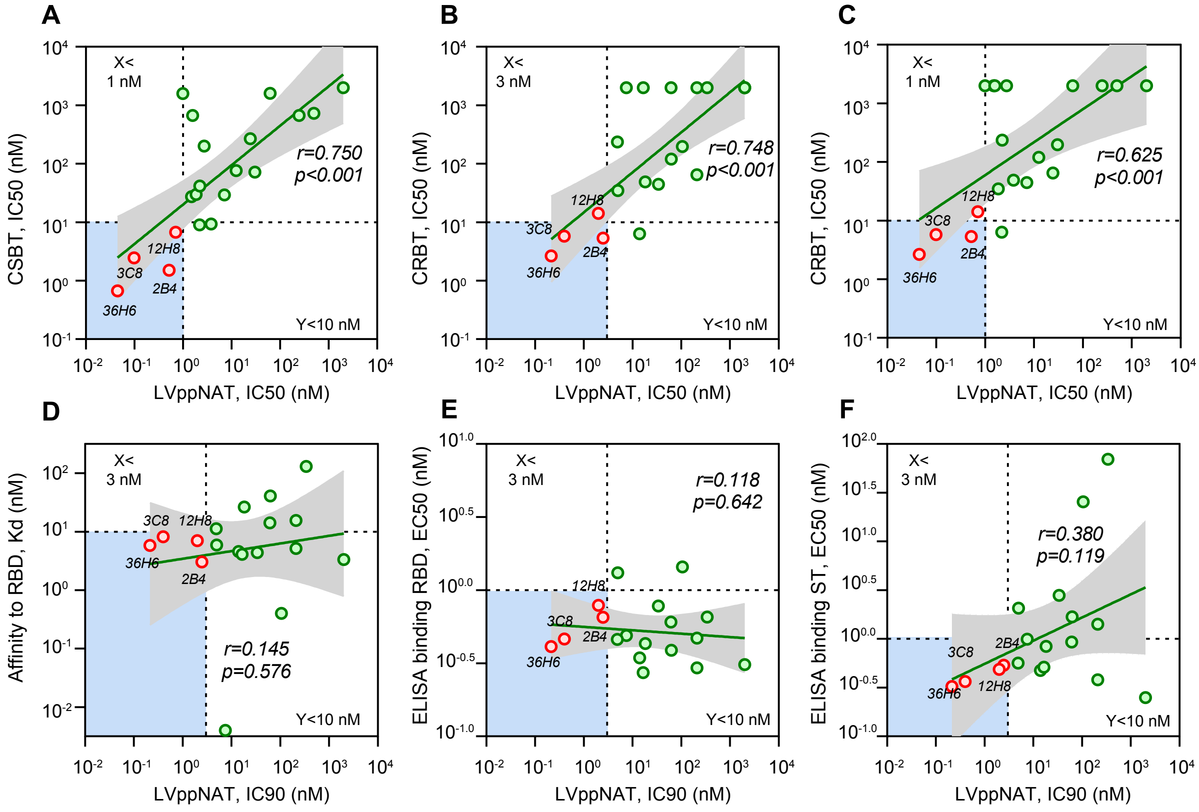

A
B
C
D
E
F
X<
1 nM
X<
3 nM
Y<10 nM
r=0.750
p<0.001
r=0.748
p<0.001
r=0.625
p<0.001
r=0.145
p=0.576
r=0.118
p=0.642
r=0.380
p=0.119
3C8
12H8
2B4
36H6
CSBT, IC50 (nM)
CRBT, IC50 (nM)
Affinity to RBD, Kd (nM)
ELISA binding RBD, EC50 (nM)
ELISA binding ST, EC50 (nM)
LVppNAT, IC50 (nM)
LVppNAT, IC90 (nM)

#### Figure S10

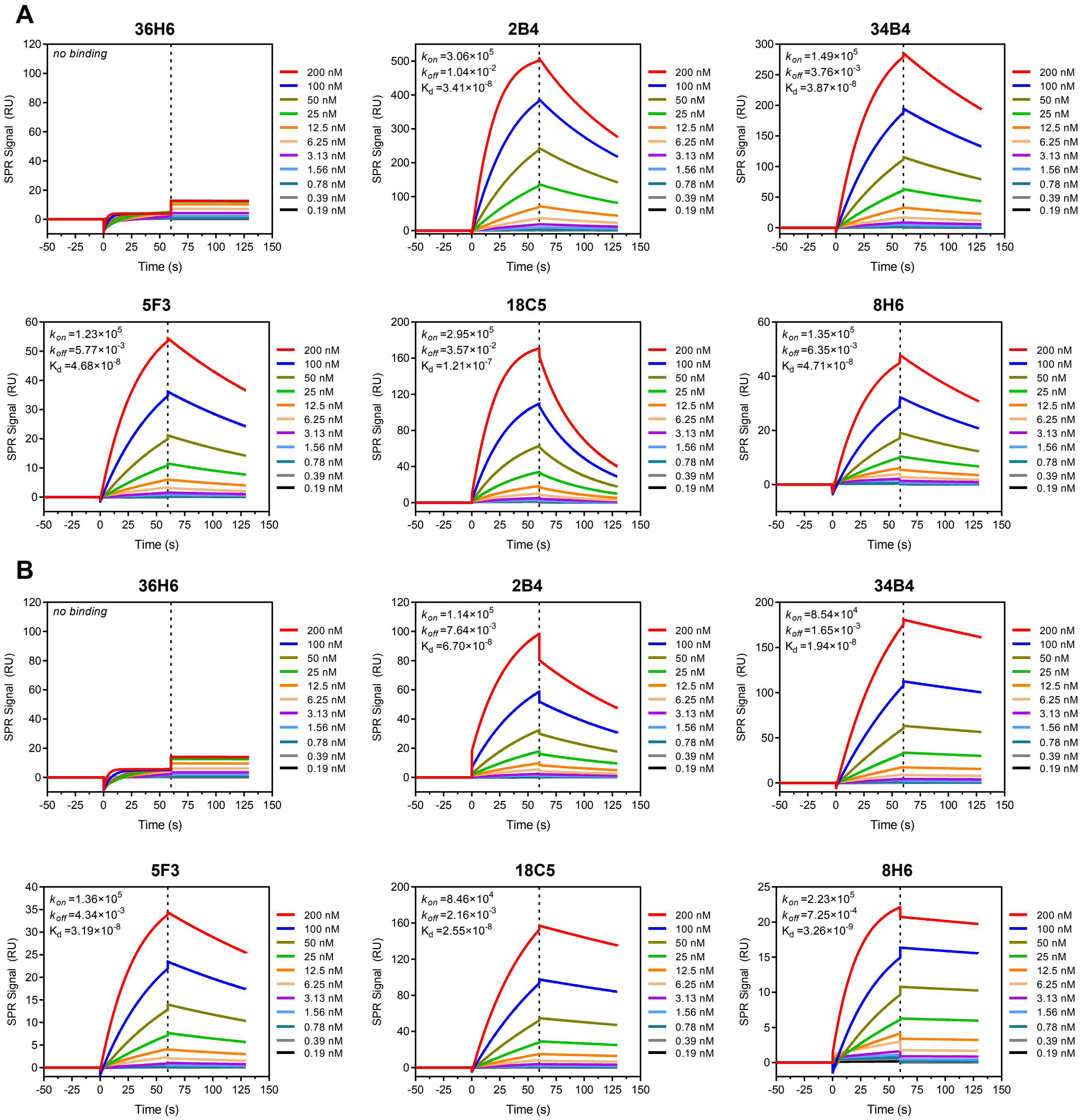
A
B
36H6
no binding
2B4
$k_{on}$ =3.06×10^5
$k_{off}$ =1.04×10^-2
$K_d$ =3.41×10^-8
34B4
$k_{on}$ =1.49×10^5
$k_{off}$ =3.76×10^-3
$K_d$ =3.87×10^-8
5F3
$k_{on}$ =1.23×10^5
$k_{off}$ =5.77×10^-3
$K_d$ =4.68×10^-8
18C5
$k_{on}$ =2.95×10^5
$k_{off}$ =3.57×10^-2
$K_d$ =1.21×10^-7
8H6
$k_{on}$ =1.35×10^5
$k_{off}$ =6.35×10^-3
$K_d$ =4.71×10^-8
36H6
no binding
2B4
$k_{on}$ =1.14×10^5
$k_{off}$ =7.64×10^-3
$K_d$ =6.70×10^-8
34B4
$k_{on}$ =8.54×10^4
$k_{off}$ =1.65×10^-3
$K_d$ =1.94×10^-8
5F3
$k_{on}$ =1.36×10^5
$k_{off}$ =4.34×10^-3
$K_d$ =3.19×10^-8
18C5
$k_{on}$ =8.46×10^4
$k_{off}$ =2.16×10^-3
$K_d$ =2.55×10^-8
8H6
$k_{on}$ =2.23×10^5
$k_{off}$ =7.25×10^-4
$K_d$ =3.26×10^-9
200 nM
100 nM
50 nM
25 nM
12.5 nM
6.25 nM
3.13 nM
1.56 nM
0.78 nM
0.39 nM
0.19 nM
SPR Signal (RU)
Time (s)

**Figure S11**

A

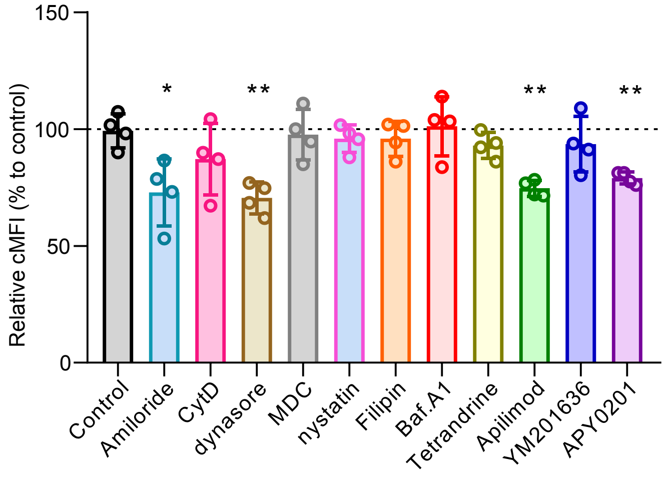

B

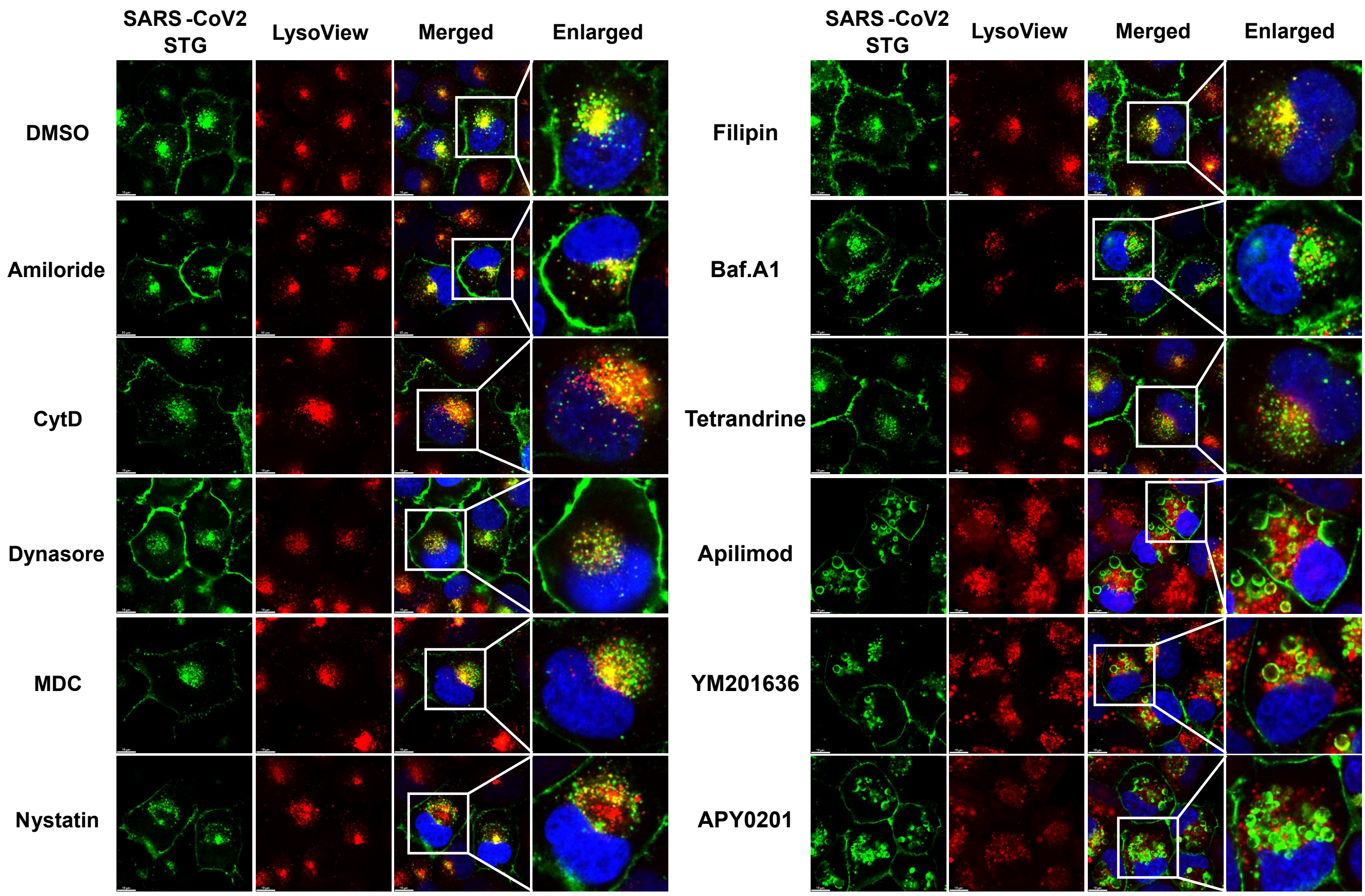

### Figure S12

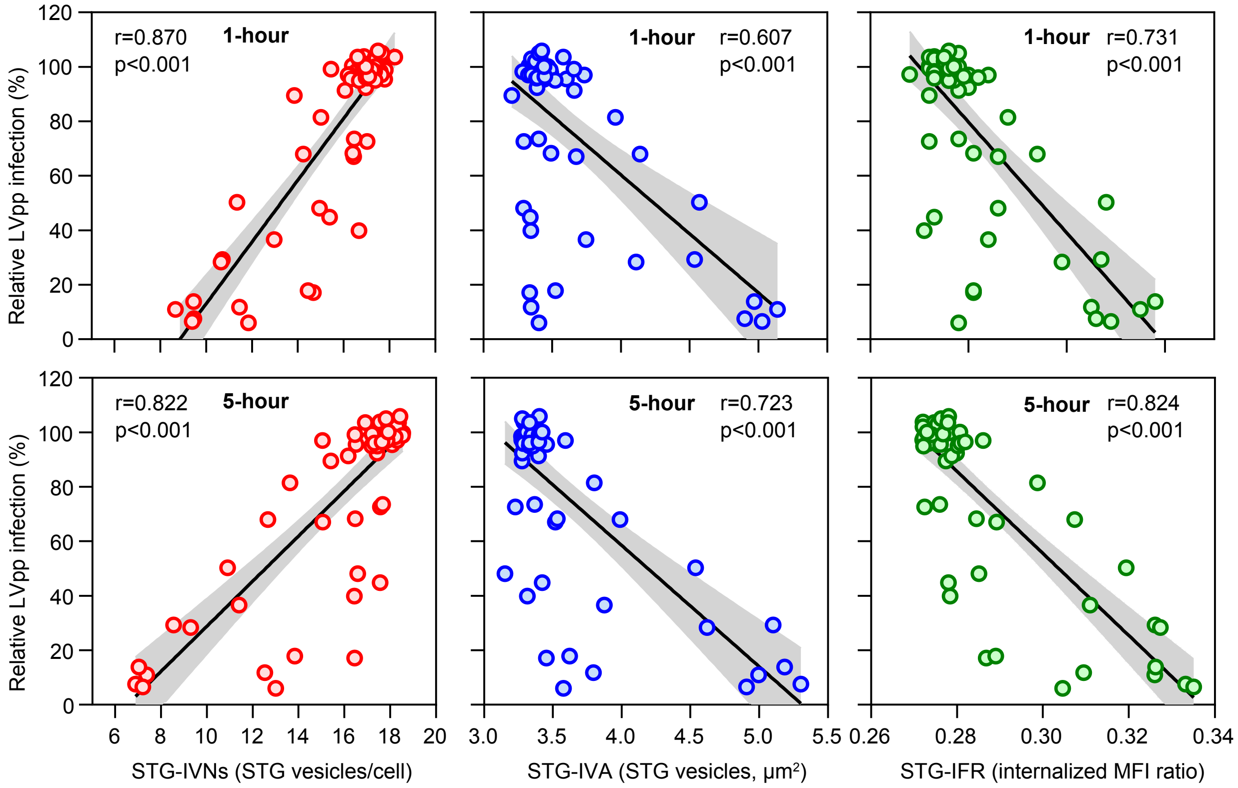

r=0.870
p<0.001
1-hour
1-hour
r=0.607
p<0.001
1-hour
r=0.731
p<0.001
r=0.822
p<0.001
5-hour
5-hour
r=0.723
p<0.001
5-hour
r=0.824
p<0.001
Relative LVpp infection (%)
Relative LVpp infection (%)
0
20
40
60
80
100
120
6
8
10
12
14
16
18
20
3.0
3.5
4.0
4.5
5.0
5.5
0.26
0.28
0.30
0.32
0.34
STG-IVNs (STG vesicles/cell)
STG-IVA (STG vesicles, µm²)
STG-IFR (internalized MFI ratio)

Table S1. Characteristics of convalescent COVID-19 patients and theirs samples involved in this study.

| | Characteristics |
|---|---|
| No. | 32 |
| Gender | |
| Male(no., %) | 19, 59.4 |
| Female(no., %) | 13, 40.6 |
| Age | |
| Median(range) | 56.5 (28-75) |
| Plasma collected at day after onset | |
| Median(range) | 33 (14-54) |
| RBD-TAb ELISA titer, $log_{10}$ SCO | |
| Mean±SD | 2.48±0.61 |
| RBD-IgG ELISA titer, $log_{10}$ SCO | |
| Mean±SD | 1.98±0.68 |
| RBD-IgM ELISA titer, $log_{10}$ SCO | |
| Mean±SD | 1.33±1.77 |
| LVppNAT, $log_{10}$ ID50 | |
| Mean±SD | 2.96±0.39 |
| CSBT titer, $log_{10}$ ID25 | |
| Mean±SD | 1.90±0.43 |
| CRBT titer, $log_{10}$ ID25 | |
| Mean±SD | 1.22±0.37 |

Table S2. Correlations between antibody titers of 32 COVID-19-convalescent human plasmas determined by various assays.

| | TAb | IgG | IgM | CRBT | CSBT | LVppNAT |
|---|---|---|---|---|---|---|
| TAb | *1.000* | | | | | |
| IgG | *0.702* | *1.000* | | | | |
| IgM | *0.584* | *0.443* | *1.000* | | | |
| CRBT* | *0.484* | *0.471* | *0.390* | *1.000* | | |
| CSBT | *0.615* | *0.504* | *0.426* | *0.645* | *1.000* | |
| LVppNAT | *0.665* | *0.674* | *0.493* | *0.480* | *0.832* | *1.000* |

The correlation coefficients (r) between titers determined by various assays were listed. Linear regression models and Pearson correlation tests were used for correlation analyses.

*The CRBT titer was cMFI inhibition at 1:20 dilution.

Table S3. The neutralizing titer of human plasma samples against authentic SARS-CoV-2 virus.

| | Authentic SARS-CoV-2 neutralization (ID100) | | |
|---|---|---|---|
| Sample ID | Test 1 | Test 2 | Average |
| p02 | 2560 | 2560 | 2560 |
| p03 | 5120 | 2560 | 3840 |
| p04 | 2560 | 2560 | 2560 |
| p14 | 640 | 640 | 640 |
| p18 | 320 | 320 | 320 |
| p24 | 640 | 320 | 480 |
| p26 | 1280 | 1280 | 1280 |
| p27 | 1280 | 640 | 960 |
| p28 | 1280 | 1280 | 1280 |
| p30 | 1280 | 1280 | 1280 |
| p31 | 160 | 80 | 120 |
| n1 | <1 | <1 | <1 |

Table S4. A summary of the characteristics of mAbs against SARS-CoV-2 involved in this study.

| mAb ID | cELISA | Biacore affinities to RBD | | | ELISA-binding assay | | Cell-based phenotypic assay | | | |
|---|---|---|---|---|---|---|---|---|---|---|
| | Epitope binning | SARS2 | SARS1 | RaTG13 | To RBD | To Strimer | LVppNAT | | CSBT | CRBT |
| | mAb cluster | Kd,nM | Kd,nM | Kd,nM | EC50,nM | EC50,nM | IC90,nM | IC50,nM | IC50,nM | IC50,nM |
| 36H6 | C1 | 5.84 | >200 | >200 | 0.411 | 0.324 | 0.214 | 0.045 | 0.671 | 2.641 |
| 2B4 | C2a | 3.05 | 34.1 | 67.0 | 0.653 | 0.536 | 2.488 | 0.520 | 1.504 | 5.329 |
| 3C8 | C3 | 8.19 | nd | nd | 0.464 | 0.366 | 0.398 | 0.099 | 2.458 | 5.765 |
| 12H8 | C4 | 6.99 | nd | nd | 0.786 | 0.488 | 1.996 | 0.710 | 6.724 | 14.12 |
| 34B4 | C2a | 4.57 | 38.7 | 19.4 | 0.344 | 0.474 | 13.98 | 2.198 | 9.008 | 6.318 |
| 60A11 | C5a | 26.4 | nd | nd | 0.432 | 0.837 | 18.28 | 3.811 | 9.359 | 48.80 |
| 53G2 | C5a | 0.004 | nd | nd | 0.489 | 0.993 | 7.492 | 1.497 | 27.26 | >2000 |
| 5F3 | C2b | 4.39 | 46.8 | 31.9 | 0.779 | 2.790 | 33.94 | 7.140 | 29.23 | 44.60 |
| 21F7 | C3 | 5.94 | nd | nd | 1.316 | 2.067 | 4.993 | 1.862 | 29.72 | 34.60 |
| 58D2 | C5a | 11.2 | nd | nd | 0.460 | 0.563 | 4.889 | 2.236 | 41.43 | 236.4 |
| 16B12 | C6 | 40.8 | nd | nd | 0.388 | 1.681 | 62.62 | 12.51 | 76.10 | 120.2 |
| 15A9 | C6 | 0.40 | nd | nd | 1.442 | 25.53 | 106.0 | 30.32 | 72.10 | 196.7 |
| 18C5 | C5b | 14.2 | 121 | 25.5 | 0.606 | 0.928 | 61.67 | 2.744 | 200.6 | >2000 |
| 14D2 | C2a | 15.6 | nd | nd | 0.469 | 1.411 | 210.1 | 24.36 | 267.8 | 64.69 |
| 23B1 | C4 | 3.34 | nd | nd | 0.310 | 0.250 | >2000 | 246.9 | 666.4 | >2000 |
| 65G9 | C5b | 5.18 | nd | nd | 0.294 | 0.380 | 210.6 | 1.586 | 671.7 | >2000 |
| 83H7 | C5b | 4.09 | nd | nd | 0.273 | 0.512 | 16.59 | 0.990 | 1587 | >2000 |
| 8H6 | C5b | 131 | 47.1 | 3.26 | 0.655 | 69.82 | 341.3 | 62.20 | 1591 | >2000 |
| CR3022 | na | nd | nd | nd | nd | nd | >2000 | 500.0 | 731.2 | >2000 |

The mAb cluster was determined according data shown in Figure S5B. na, not applicable; nd, not detected. The binding affinities of these mAb to RBD proteins were summarized from the data shown in Figure S6 and Figure S10. The ELISA binding activities of these mAbs were summarized from the data shown in Figure S5A. The LVppNAT data were based on Figure S7C, the CSBT data were based Figure 4B, and the CRBT data were based on Figure S7A and S7B.
